# *S. pombe* DNA translocases Rrp1 and Rrp2 have distinct roles at centromeres and telomeres that ensure genome stability

**DOI:** 10.1101/738435

**Authors:** Anna Barg-Wojas, Kamila Schirmeisen, Jakub Muraszko, Karol Kramarz, Gabriela Baranowska, Antony M. Carr, Dorota Dziadkowiec

## Abstract

Homologous recombination (HR) is a DNA repair mechanism that ensures, together with heterochromatin machinery, the proper replication, structure and function of telomeres and centromeres that is essential for the maintenance of genome integrity. *Schizosaccharomyces pombe* Rrp1 and Rrp2 participate in HR and are orthologues of *Saccharomyces cerevisiae* Uls1, a SWI2/SNF2 DNA translocase and SUMO-Targeted Ubiquitin Ligase. We show that Rrp1 or Rrp2 upregulation leads to chromosome instability and growth defects. These phenotypes depend on putative DNA translocase activities of Rrp1 and Rrp2. Either Rrp1 or Rrp2 overproduction results in a reduction in global histone levels, suggesting that Rrp1 and Rrp2 may modulate nucleosome dynamics. In addition we show that Rrp2, but not Rrp1, acts at telomeres. We propose that this role depends on the previously described interaction between Rrp2 and Top2. We conclude that Rrp1 and Rrp2 have important roles for centromere and telomere function and maintenance, contributing to the preservation of genome stability during vegetative cell growth.

**SUMMARY STATEMENT:** *Schizosaccharomyces pombe* DNA translocases Rrp1 and Rrp2 modulate centromere and telomere maintenance pathways and dysregulation of their activity leads to genome instability.

## INTRODUCTION

Homologous recombination (HR) is a highly conserved pathway that functions during DNA replication, participates in the repair of double strand breaks (DSBs) during interphase and is essential for meiosis. During HR the key Rad51 recombinase forms a nucleofilament on single stranded DNA which catalyses strand invasion into intact homologous double stranded DNA (Symington, 2002). Rad51 is aided by a group of proteins called recombination “mediators”.

In *S. pombe* two mediator complexes have been shown to act in parallel to promote Rad51-dependent strand exchange: Rad55-Rad57 and Sfr1-Swi5 (Akamatsu et al., 2007), both of which are conserved in humans (Yuan and Chen, 2011). We previously identified another complex, Rrp1-Rrp2, that acts in a Swi5/Sfr1-dependent sub-pathway of HR in the replication stress response (Dziadkowiec et al., 2009). Both Rrp1 and Rrp2 are orthologues of *S. cerevisiae* Uls1, a SWI2/SNF2 DNA translocase and SUMO-Targeted Ubiquitin Ligase (STUbL). We subsequently demonstrated that Rrp1-Rrp2 can function to negatively regulate one or more sub-pathways of Rad51-mediated recombination (Dziadkowiec et al., 2013).

Telomeres and centromeres are potentially difficult to replicate regions due to the presence of repetitive sequences that can form secondary structures impeding replication fork progression. These repetitive sequences are often unstable and constitute the hotspots of replication fork arrest and recombination. HR proteins act at arrested replication forks: Rad51 binding promotes the stability of the fork itself (Mizuno et al., 2013; Schlacher et al., 2011) whereas when the fork is inactivated, the strand exchange activity of Rad51 promotes the reconstitution of replication (Lambert et al., 2005; McGlynn and Lloyd, 2002). In *S. pombe*, the telomere binding protein Taz1 (and in mammals its orthologue TRF1) attenuates the tendency of telomere repeats to block replication (Miller et al., 2005; Sfeir et al., 2009). In *taz1*Δ mutants, arrested replication forks at telomeres are incorrectly processed, which leads to telomeric entanglements that cannot be resolved at temperatures below 20°C. The consequence of this is the formation of chromosome bridges, chromosome missegregation and reduced cell viability (Miller and Cooper, 2003).

Topoisomerase II (Top2) is essential for telomere maintenance and mutants such as *top2-191*, characterized by slower catalytic turnover, are able to supress *taz1*Δ telomeric entanglement phenotypes (Germe et al., 2009). It has recently been shown that Rrp2 protects cells from Top2-induced DNA damage and its absence is toxic in *top2-191* mutant (Wei et al. 2017) suggesting a role for Rrp2 in telomere replication. This function is shared with its *S. cerevisiae* orthologue, Uls1, which has been shown to inhibit nonhomologous end joining at telomeres (Lescasse et al., 2013) and to protect the cells against Top2 poisons in a manner dependent on its ATPase activity and SUMO-binding (Wei et al., 2017).

Fission yeast centromeres are composed of large tandem and inverted repeats (*dg* and *dh*) surrounding a unique central core where histone H3 is replaced by the histone variant CENP-A (Cnp1) to allow kinetochore formation. *dg* and *dh* elements are assembled into heterochromatin that both imposes transcriptional silencing at this region and is required for accurate chromosome segregation (Allshire and Ekwall, 2014; Ekwall, 2007). Several trans-acting factors are required for heterochromatin formation, the most important being Clr4, which methylates histone H3 on lysine 9. This methylation results in the binding of Swi6, the *S. pombe* HP1 homolog (Ekwall et al., 1995; Nakayama et al., 2001). Interestingly, Rad51 recombinase localizes to the centromere and has a role in the suppression of rearrangements within its repeats (Nakamura et al., 2008).

It was recently reported (Onaka et al., 2016) that depletion of Rad51 also impairs transcriptional repression of genes inserted within centromeres and results in elevated levels of chromosome loss. This indicates that, even though the centromeres are largely assembled into heterochromatin, HR factors not only promote recombination between centromere repeats but are also important for the proper formation and/or function of heterochromatin. Consistent with these observations, Rad51 and Rad52 have also been shown to regulate CENP-A (CaCse4) Ievels in *Candida albicans* centromeres (Mitra et al., 2014) and, in *S. pombe*, the Rqh1-Top3 helicase-topoisomerase complex influences centromere topology by regulating CENP-A (Cnp1) levels at the core region (Norman-Axelsson et al., 2013). This function of Rqh1-Top3 is independent from its role in Holiday junction dissolution. Rqh1 is a member of the RecQ family of helicases that includes the human WRN and BLM proteins. RecQ-family helicases are involved in the processing of stalled replication forks and have been reported to play a role at telomeres when replication fork progression is perturbed (Barefield and Karlseder, 2012; Rog et al., 2009). The regulation of centromere and telomere integrity by proteins involved in HR and the replication stress response is clearly complex and the characterisation of additional factors participating in this process is therefore crucial for a better understanding of centromere and telomere biology.

In this report we show that Rrp1 and Rrp2 are differentially involved in the maintenance of centromere function (assayed as TBZ sensitivity) and structure (assayed as disruption of centromeric silencing). Both effects are especially evident in cells where centromeres are destabilized by the loss of the heterochromatin proteins Clr4 and Swi6. The role of Rrp1 at centromeres is more pronounced than that of Rrp2, depends on a functional ATPase domain and, to a lesser extent, on an intact RING domain. Rrp2 activity at the centromere mostly depends on functional RING and ATPase domain, with the SIM motifs playing a less important role. We also show that Rrp2 has a separate function at telomeres. This requires the Rrp2 SUMO binding and ATP binding domains, but not the RING domain. Additionally, our data suggest that Rrp1, similarly to what has been shown for Rrp2 (Wei et al., 2017), may have ATP-dependent translocase activity. We propose that both Rrp1 and Rrp2 proteins have important and non-redundant roles in the maintenance of repetitive genomic regions and that dysregulation of their activity leads to genetic instability.

## RESULTS

### Rrp1 and Rrp2 may have a role in centromere function

Rrp1 and Rrp2 are involved in replication stress responses in a Swi5-Sfr1 dependent branch of homologous recombination (Dziadkowiec et al., 2009; Dziadkowiec et al., 2013). Previous work has demonstrated that HR factors such as Rad51 and Rad54 are required to ensure centromere stability and that cells devoid of these proteins are sensitive to thiabendazole (TBZ), a microtubule-destabilizing agent (Onaka et al., 2016). We also find that a *rad57*Δ mutant is sensitive to TBZ, but that *swi5*Δ, *sfr1*Δ, *rrp1*Δ and *rrp2*Δ mutants are not. A small increase in TBZ sensitivity is observed in *rad57*Δ*rrp1*Δ and *rad57*Δ*rrp2*Δ strains when compared to *rad57*Δ (Fig. S1A), implying that Rrp1 and Rrp2 may have a role in centromere maintenance. Previous work (Li et al., 2013) has shown that that replication fork stability is required, together with heterochromatin, to ensure centromere integrity. We thus reasoned that, in mutants sensitive to TBZ due to destabilization of heterochromatin at centromeres (Allshire et al., 1995; Ekwall et al., 1996), this effect would be more pronounced. Unexpectedly, we observed that *rrp1*+ or *rrp2*+ deletion slightly decreased TBZ sensitivity in *swi6*Δ background, and in the *clr4*Δ mutant background deletion of *rrp1*+, but not of *rrp2*+, lead to the rescue of the growth defect (Fig. S1B). This suggests that Rrp1 (and possibly Rrp2 also) contributes to centromere maintenance and, when heterochromatin structure is disrupted, their activity is deleterious.

In this context we reasoned that overproduction of Rrp1 and Rrp2 should result in growth defect and/or TBZ sensitivity in *swi6*Δ and *clr4*Δ mutants and that this may also be apparent in wild type cells. We thus examined the effect of *rrp1*+ or *rrp2*+ over-expression from the medium strength *nmt* promoter (*nmt41* and *42*). Indeed, this caused viability loss and moderate TBZ sensitivity in otherwise wild type cells grown under unperturbed conditions (Fig. 1A see also Fig. S4C). The growth defect induced by either *rrp1*+ (Fig. 1B) or *rrp2*+ (Fig. 1C) over-expression in the *swi6*Δ mutant was exacerbated when compared to that seen in wild type cells and their over-expression strongly sensitized *swi6*Δ cells to TBZ. Furthermore, the viability loss caused by *rrp1*+ and *rrp2*+ over-expression in the *clr4*Δ mutant was greater than in WT or *swi6*Δ cells (Fig. 1B,C). This induced us to propose a hypothesis that Rrp1 and Rrp2 can affect centromere function and the effect of copy number dysregulation becomes more pronounced as centromeres become more dysfunctional.

**Figure 1.**
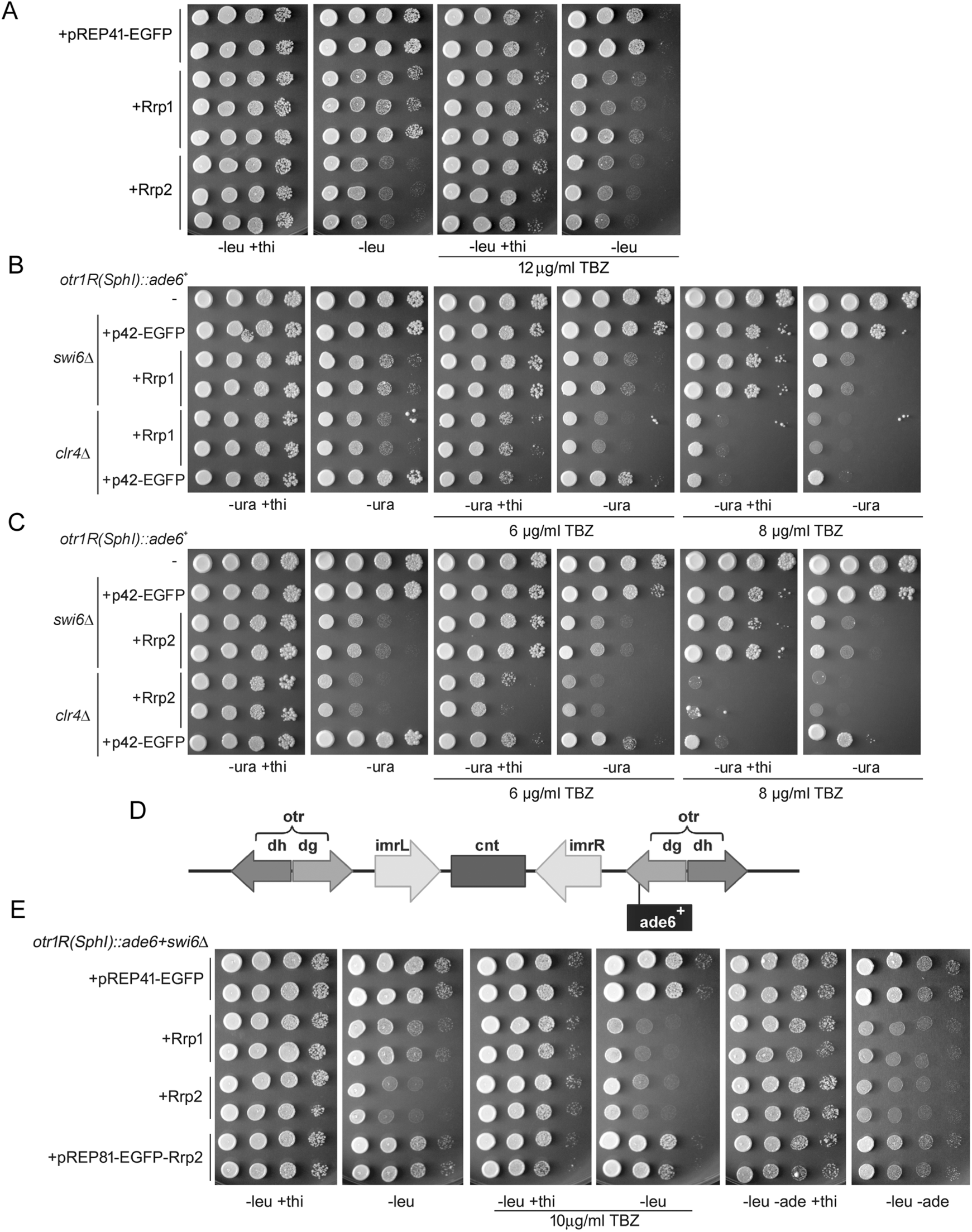
Up regulation of Rrp1 and Rrp2 level causes genetic instability. (A) Induction of *rrp1*+ or *rrp2*+ expression leads to growth defect and increase in TBZ sensitivity. Rrp1 and Rrp2 act independently of Swi6 and Clr4 as over-expression of (B) *rrp1*+ and (C) *rrp2*+ sensitizes *swi6*Δ and *clr4*Δ cells to TBZ. (D) The diagram depicting the localization of the *ade6*+ gene in the *dg* region of centromere 1 in a strain (*otr1R(SphI)::ade6*+) used to monitor centromere silencing. (E) Over-expression of *rrp1*+ and *rrp2*+ reverses the transcriptional de-repression of the *dg*-*ade6*+ gene that is conferred by *swi6*Δ.

Loss of transcriptional silencing at the centromere is a further indicator of the disruption of centromere structure (Allshire et al., 1995). We thus examined the effect of Rrp1 and Rrp2 on silencing by assaying functional expression of an *ade6*+ gene that is inserted in the *dg* region of centromere 1 (*dg*-*ade6*+: Fig. 1D). Interestingly, we observed that over-expression of *rrp1*+ or *rrp2*+ increased the repression of the *dg*-*ade6*+ gene in an otherwise wild type strain, as determined by the lack of growth on plates devoid of adenine (Fig. S2A). This phenotype is mild, so we also examined the effect of *rrp1*+ or *rrp2*+ in a *swi6*Δ mutant where silencing is alleviated. Intriguingly, we observed that *rrp1*+ over-expression reversed the transcriptional de-repression of *dg*-*ade6*+ in *swi6*Δ cells (Fig. 1E). The strong viability loss conferred by *nmt41*-driven *rrp2*+ over-expression made it difficult to assess its role in silencing. However, when *rrp2*+ is expressed from the lower-strength *nmt81* promoter (which leads to growth inhibition comparable to that caused by *nmt41*-*rrp1*+ over-expression), we observed a modest but reproducible increase in the repression of *dg*-*ade6*+ and TBZ sensitivity, albeit less significant than that seen for *nmt41*-*rrp1*+ (Fig. 1E). The difference in *rrp1*+ or *rrp2*+ over-expression phenotypes will be discussed later.

These results support our hypothesis that even though deletion of *rrp1*+ or *rrp2*+ has no effect on the silencing of *dg*-*ade6*+ in *swi6*+ or *swi6*Δ backgrounds (Fig. S2B) Rrp1 and Rrp2 may have roles that are important for proper centromere function that become particularly apparent in mutants, such as *swi6*Δ and *clr4*Δ, where centromere structure is already disrupted.

### Increase in Rrp1 and Rrp2 copy number results in chromosome instability

It has been proposed (Javerzat et al., 1996) that proteins involved in the formation of centromere/kinetochore complexes should be present in cells in precise quantities to achieve correct assembly of these structures. Dysregulation of gene copy numbers for such proteins would thus result in elevated levels of chromosome instability. Microscopic examination of DAPI stained cells revealed that prolonged over-expression of *rrp1*+, and even more so of *rrp2*+, causes mitotic aberrations including chromosome non-disjunction and “cut” nuclei (Fig. 2A). We observe in these cells the appearance of bright clusters of Rad11 (RPA) foci (Fig. 2B), similar to those seen when replication was purturbed in the absence of γH2AX (Mejia-Ramirez et al., 2015). Additionally, approximatelty 30 % of anaphase cells over-expressing *rrp1*+ or *rrp2*+ accumulate fragmented DNA and bridges coated with Rad11 (inset with arrowhead in Fig. 2B). This implies that chromosome segregation defects occur in these cells.

**Figure 2.**
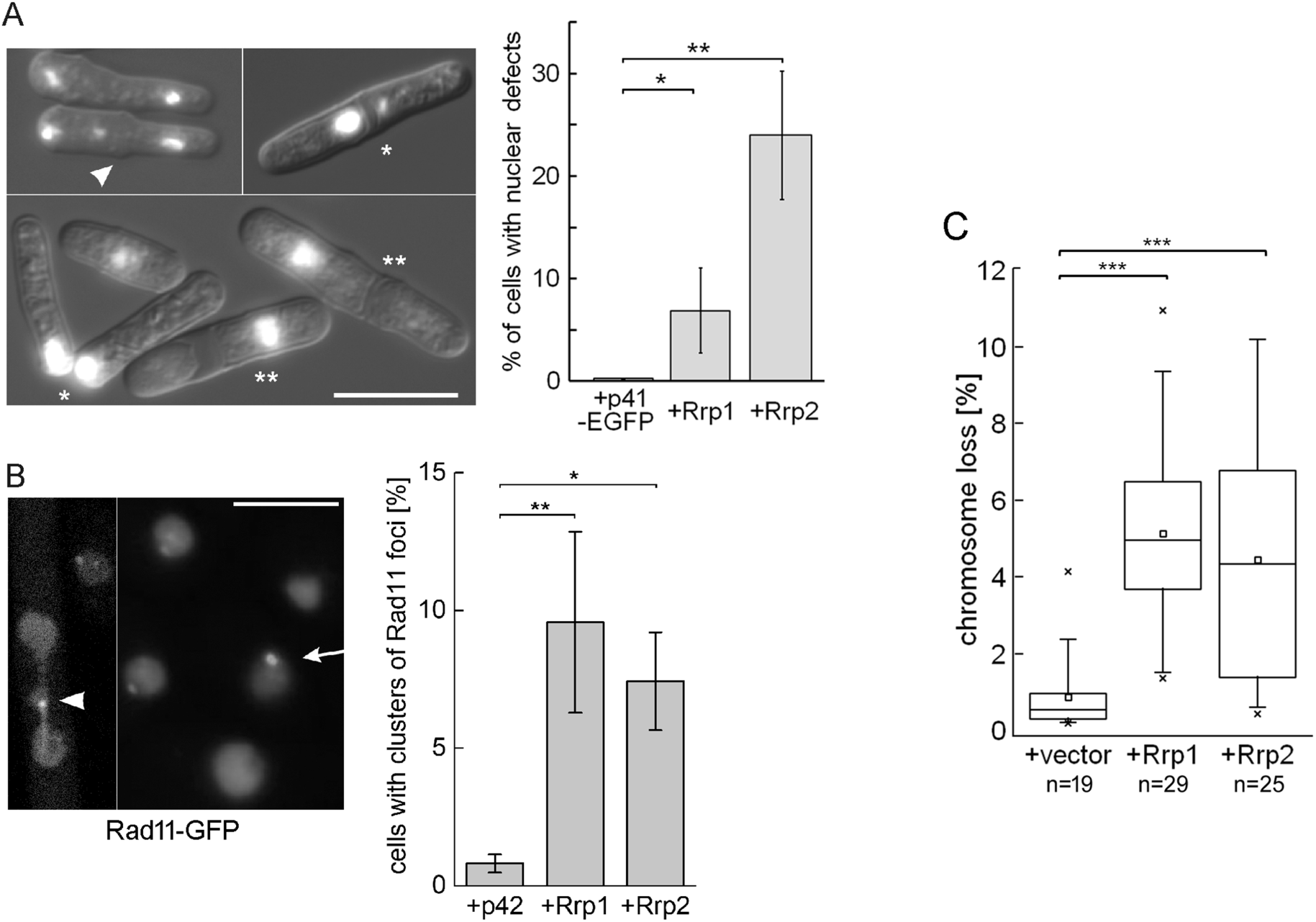
Over-expression of *rrp1*+ and *rrp2*+ impairs DNA segregation. Induction of *rrp1*+ or *rrp2*+ expression leads to (A) mitotic aberrations, such as lagging/stretched chromosomes/anaphase bridges (marked as white arrowhead), cut (marked as *) and non-disjunction (marked as **) as observed by DAPI staining of the nuclei. 5 independent transformants for vector, *rrp1*+ or *rrp2*+ were analysed and the total number of cells counted was above 1000. (B) Bright clusters of Rad11 foci (marked as white arrow) and Rad11 coated DNA bridges (marked as white arrowhead) accumulate in *rrp1*+ or *rrp2*+ over-expressing cells. Regular Rad11 foci are unmarked. The experiment was repeated 3 times and the total number of cells counted was above 2000. Scale bar represents 10 µm. (C) Induction of *rrp1*+ or *rrp2*+ expression leads to the loss of the nonessential Ch^16^ minichromosome carrying the *ade6*-*216* allele, resulting in red colony formation on medium with limiting adenine concentration. Data from two independent transformations (n= number of colonies) were analysed.

Problems associated with chromosome segregation result in chromosome instability (Murray et al., 1994). We used a strain with a nonessential Ch^16^ minichromosome carrying the *ade6*-*216* allele *trans-*complementing the endogenous *ade6*-*210* allele of the host cell to measure chromosome loss induced by *rrp1*+ and *rrp2*+ over-expression. In this assay cells that lose the minichromosome form red colonies when grown on medium with a limiting concentration of adenine. We show that *rrp1*+ or *rrp2*+ over-expression increased minichromosome loss 5-fold (Fig. 2C).

When considered together, these data (Figs. 1-2 and Fig. S1) indicate that dysregulation of Rrp1 or Rrp2 interferes with centromere function, resulting in chromosome segregation defects and genetic instability.

### Interdependence of *rrp1*+ and *rrp2*+ over-expression phenotypes

Previous genetic analysis of *rrp1*Δ and *rrp2*Δ cells in response to DNA damage implied that these two proteins function as a unit (Dziadkowiec et al., 2009). Here we observed differences between the two gene functions: upon *rrp2*+ overexpression the growth defect is greater and the TBZ sensitivity and transcriptional repression are lower when compared to *rrp1*+ over-expressing cells (Fig. 1E). Importantly, the growth defect, increased TBZ sensitivity and silencing caused by over-expression of *rrp1*+ or *rrp2*+ are not dependent on the presence of their respective paralogue (Fig. S3A). When both genes are over-expressed, the resulting growth defect is equivalent to that caused by over-expression of *rrp2*+ (Fig. S3B). Furthermore, the increased toxicity of *rrp2*+ as compared with *rrp1*+ over-expression is partially dependent on the presence of Rad51 recombinase (Fig. S3C). These data suggest the existence of separate roles for both proteins and imply that Rrp2 toxicity might also result from its activity at other loci than the centromere. Accordingly, recent work demonstrated an Rrp1-independent function for Rrp2 in regulating Top2 degradation (Wei et al., 2017). Thus, Rrp1 and Rrp2 have activities that are independent of the other paralogue.

### Rrp1 and Rrp2 can bind to centromeric and/or telomeric regions

We have previously shown that Rrp1 and Rrp2 form foci co-localizing with MMS-induced Rad52 foci at sites of DNA damage (Dziadkowiec et al., 2009; Dziadkowiec et al., 2013). Upon prolonged over-expression EGFP-tagged Rrp1 and Rrp2 bind to DNA and also form spontaneous foci in cells. Co-staining for EGFP-Rrp1 or EGFP-Rrp2 with ECFP-Swi6 demonstrated that >40% of Rrp foci are formed in perinuclear regions and co-localize with Swi6 foci (Fig. 3A) suggesting that Rrp1 and Rrp2 can bind to centromeres and/or telomeres. However, Rrp1 and Rrp2 foci are formed in the absence of Swi6 and Clr4 (Fig. 3B) so Rrp1 and Rrp2 localization is not dependent on heterochromatin. Interestingly, we observe that about 70% of Rrp1 and Rrp2 foci co-localize with Rad11 (Fig. 3C). We thus propose that Rrp1 and Rrp2 bind to sites of accumulation of DNA damage and/or ssDNA throughout the genome but that the detrimental effect of their dysregulation is most pronounced at centromeres and/or telomeres.

**Figure 3.**
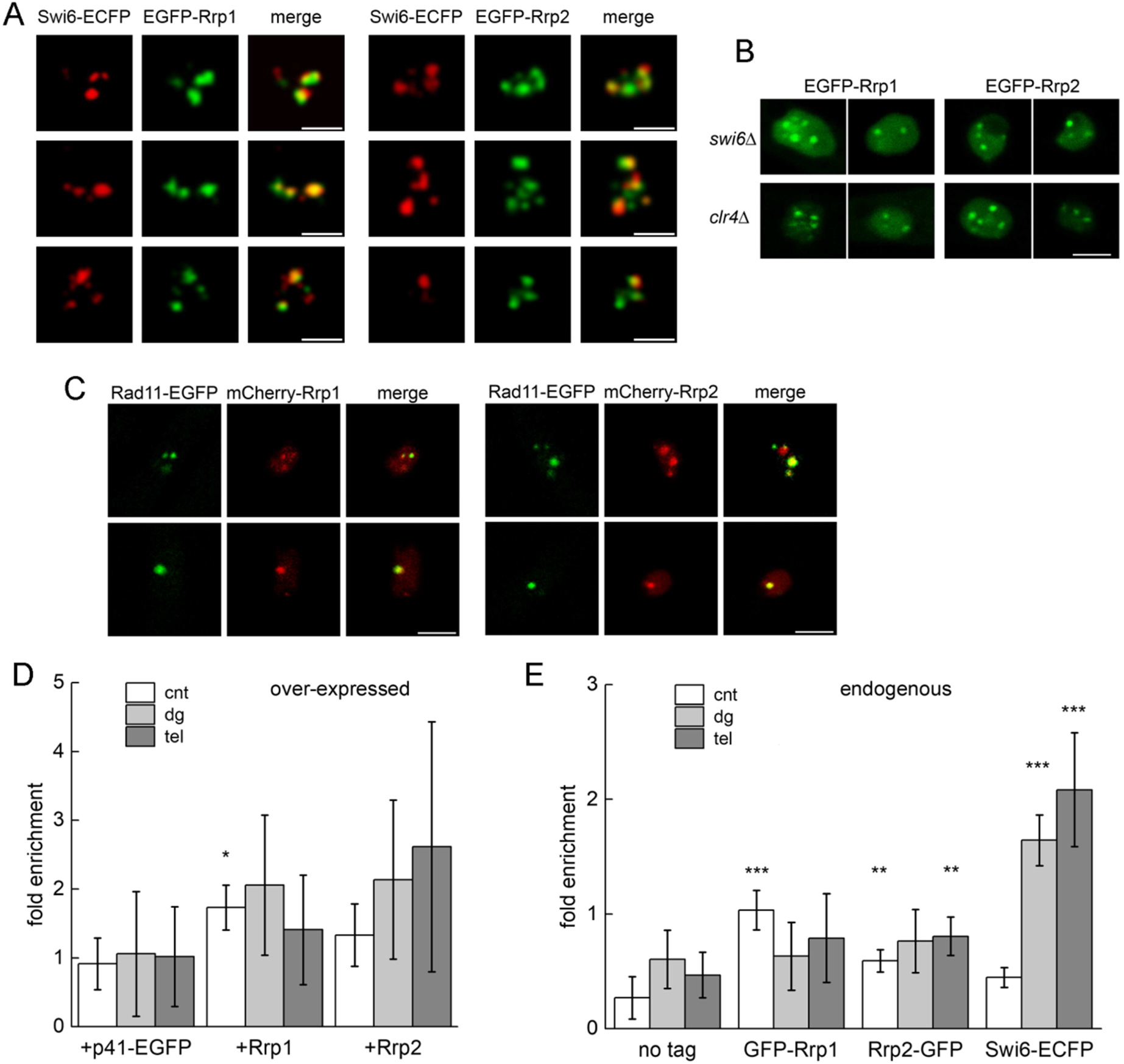
Rrp1 and Rrp2 can bind to centromeric and telomeric DNA. (A) Rrp1 and Rrp2 nuclear foci co-localize in about 40% with foci for Swi6. Strain expressing Swi6-ECFP from endogenous loci was transformed with pREP41 plasmids carrying genes for Rrp1-EGFP or Rrp2-EGFP. Two independent transformants were analysed for each assay and at least 100 cells positive for both ECFP and EGFP signal were counted. Scale bar represents 2 µm. (B) Rrp1 and Rrp2 foci formation is independent from the presence of Swi6 and Clr4. *swi6*Δ and *clr4*Δ strains over-expressing *EGFP-rrp1*+, *EGFP-rrp2*+ were examined. Scale bar represents 10 µm. (C) Numerous Rrp1 and Rrp2 nuclear foci colocalize with foci for Rad11. Strain expressing Rad11-GFP from endogenous loci was transformed with pREP41 plasmids carrying genes for Rrp1-mCherry or Rrp2-mCherry. Scale bar represents 2 µm. (D) Over-expressed and (E) endogenous Rrp1 and Rrp2 can bind to centromere and/or telomere region as shown by chromatin immunoprecipitation. Fold enrichment is calculated relative to actin control. Primers were located in centromere core (cnt), outer repeat (dg) and telomere (tel) regions. Strain expressing endogenous Swi6-ECFP was used as a positive control. Data for over-expressed proteins were obtained from two and for endogenous from four independent experiments, for each, real time PCR was repeated twice.

Accordingly, ChIP of over-expressed EGFP-tagged Rrp1 and Rrp2 indicated that both proteins associated preferentially with centromere and telomere. However, as Rrp1 and Rrp2 are also likely to associate with chromatin across the genome, their enrichment at these loci over actin control is not strong (Fig. 3D). While there are only an average of 1-2 molecules of RNA for Rrp1 and Rrp2 per cell (Quantitative gene expression, PomBase) and we are unable to detect these proteins when tagged at their native locus, ChIP of endogenously GFP-tagged Rrp1 and Rrp2 was also consistent with an ability of the endogenous proteins to associate with centromeres and telomeres (Fig. 3E).

### Disruption of centromere function conferred by Rrp1 and Rrp2 overproduction depends differentially on their domains

Rrp1 and Rrp2 have a complex domain structure (Dziadkowiec et al., 2009) (Fig. 4A). We examined the importance of these domains for the induced growth defect, TBZ sensitivity and for silencing of *dg*-*ade6*+ in the *swi6*Δ background. We first confirmed that Rrp1 and Rrp2 with either Walker-B mutations (Rrp1-DAEA, Rrp2-DAEA), RING mutations (Rrp1-CS, Rrp2-CS) and Rrp2 with the 6 SIMs mutated (Rrp2-SIM) were expressed (Fig. S4A). All the mutant proteins form foci in the nucleus (Fig. S4B), albeit for Rrp1-DAEA their number is somewhat decreased. All domains contribute to the loss of viability conferred by *rrp1*+ or *rrp2*+ over-expression (Fig. S4C).

**Figure 4.**
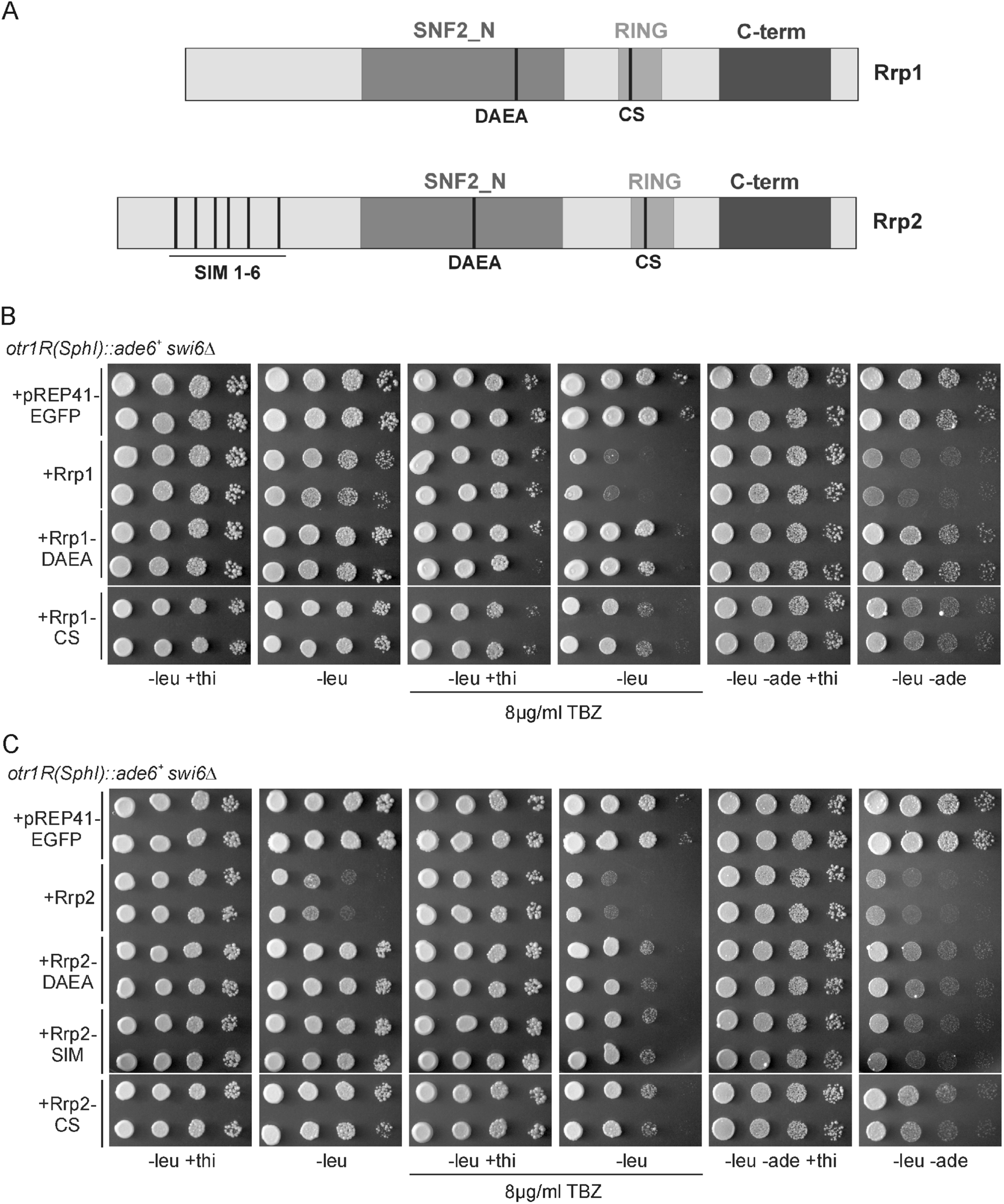
Translocase activity and RING domain of Rrp1 and Rrp2 have distinct roles in the maintenance of centromere. Rrp1 and Rrp2 have a complex domain structure and (A) mutations abolishing their putative SWI2/SNF2 DNA translocase (DAEA) and ubiquitin ligase (CS) activity and SUMO binding (SIM) are shown. (B) Functional Rrp1 translocase and an intact RING domain are required for toxicity, TBZ sensitivity and the silencing of *dg*-*ade6*+ gene, whereas for Rrp2 (C) all activities are important for toxicity and TBZ sensitivity but SIM motifs are not required for silencing of *dg*-*ade6*+ gene. Cells were transformed with plasmids harbouring genes for wild type or mutated versions of respective proteins and the ability of the constructs to support growth on plates with TBZ, as well as to repress growth on plates without adenine under conditions inducing gene expression was assessed. TBZ was added to the plates at the indicated concentrations.

Over-expression of *rrp1-DAEA* or *rrp1-CS* does not cause marked growth defects or TBZ sensitivity in the *swi6*Δ background (Fig. 4B), although only Rrp1-DAEA completely loses its toxicity. Similarly, over-expression of *rrp1-DAEA* fails to increase transcriptional repression of *dg*-*ade6*+, while *rrp1-CS* shows an intermediate phenotype (Fig. 4B). This demonstrates that the putative translocase activity of Rrp1 is critical both for growth inhibition and its role at the centromere and that the RING domain is involved, but less important.

However, when we examined the role of *rrp2* mutants the situation was different. Over-expression of all mutated *rrp2* genes in the *swi6*Δ background does not cause marked growth defect and leads to similar, attenuated TBZ sensitivity (Fig. 4C). In contrast to *rrp1* however, over-expression of *rrp2-DAEA* and *rrp2-CS* results in similar, intermediate levels of TBZ sensitivity and *dg-ade6*+ silencing, demonstrating that these domains are involved in, but not critical for, proper centromere structure and function. What’s more, SIM domains, while important for growth defect and TBZ sensitivity, are not significantly influencing transcriptional repression (Fig. 4C). This indicates that, for Rrp1, its overexpression-induced growth defect and TBZ sensitivity are closely related to disruption of centromere structure and depend mostly on its putative translocase activity. However, the growth defect, and to some extent the TBZ sensitivity, induced by Rrp2 over-expression are not entirely related to disruption of centromere structure and differentially depend on separate Rrp2 activities.

### Rrp1 and Rrp2 influence centromere structure by modulation of histone levels

The fact that the RING domains of Rrp1 and Rrp2 play a role in overproduction-induced toxicity (Fig. 4B,C) suggests these proteins may possess ubiquitin ligase activity. Indeed, we immunoprecipitated ubiquitylated proteins with both Rrp1 and Rrp2 (Fig. S5A) and *rrp1*+ over-expression and, to a lesser degree, *rrp2*+ over-expression leads to an accumulation of ubiquitin modified proteins (Fig. S5B). Interestingly, the ubiquitin-conjugating enzymes Rhp6, UbcP3(Ubc7) and Ubc15 have previously been identified in screens for genes whose over-expression disrupts silencing (Choi et al., 2002; Nielsen et al., 2002). Rhp6 was subsequently shown to ubiquitylate histone H2B on lysine 119 (H2B-K119). *rhp6*+ deletion, leading to loss of H2B ubiquitination, or an *H2B-K119R* mutation, both result in an increase in transcriptional repression at centromeres that is accompanied by defects in cell growth and nuclear structure (Tanny et al., 2007; Zofall and Grewal, 2007).

We thus examined if the *H2B-K119R* mutation led to the reversion of transcriptional de-repression of *dg-ade6*+ in the *swi6*Δ background and assayed for TBZ sensitivity. Indeed, the *H2B-K119-R* mutation causes transcriptional silencing of *dg-ade6*+, a marked increase in TBZ sensitivity in *swi6*Δ cells and a decrease in viability (Fig. S5C), reminiscent of the effect of *rrp1*+ or *rrp2*+ over-expression (Fig. 1E). We also observe a decrease of the levels of ubiquitylated H2B after *rrp1*+ over-expression, for *rrp2*+ the effect is not statistically significant (Fig. S5D,E). Surprisingly, we noticed that total levels of histone H2B are reduced after *rrp1*+ or *rrp2*+ over-expression (Fig. S5D,F). Thus, the ratio of H2BUbi to H2B is not changed under these conditions (Fig. S5G). This indicates that defect in H2B ubiquitylation is unlikely to be the main cause of centromere destabilization induced by *rrp1*+ or *rrp2*+ over-expression.

Interestingly, the reduction of histone levels we observe after *rrp1*+ or *rrp2*+ over-expression is not specific to H2B, as we also see a similar decrease for histone H3 (Fig. 5A,B). We therefore examined if this effect is dependent on ATPase and RING domains of Rrp1 and Rrp2, which are important for overproduction-induced toxicity of these proteins. We observed that only the potential translocase activity is required for Rrp1-induced H3 depletion, whereas both translocase and ubiquitin ligase activities are required for Rrp2-induced H3 depletion (Fig. 5A,B). This is especially evident when the intensity of the H3 signal is normalized to the intensity of GFP signal (tag on Rrp1 and Rrp2 proteins) (Fig. 5C). This demonstrates that the differences in the amount of histone H3 we observe do not stem from the differences in wild type and mutant protein levels in transformants examined and suggests that the increase in Rrp1 and Rrp2 copy numbers destabilizes nucleosomes.

**Figure 5.**
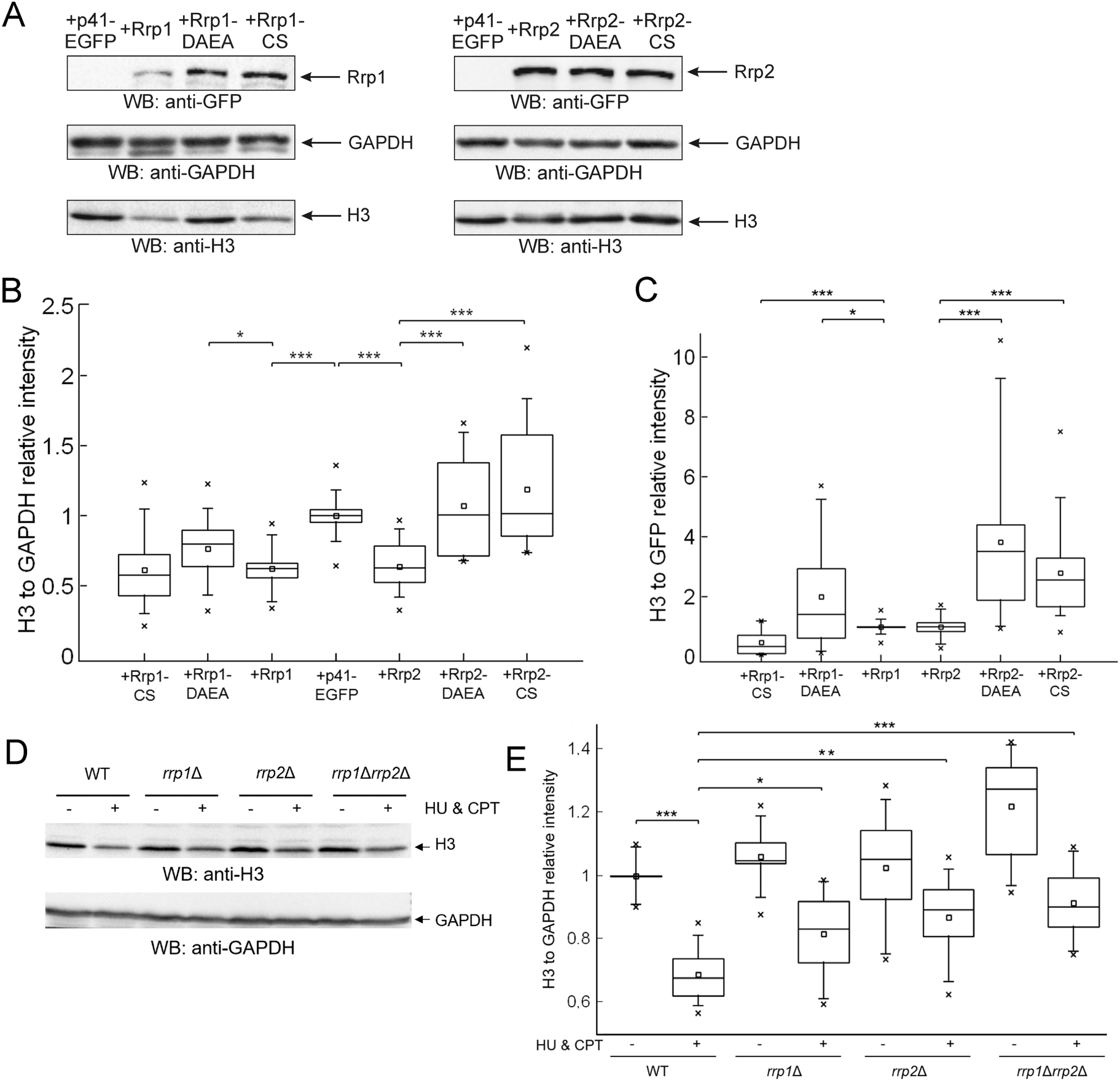
Rrp1 and Rrp2 role in centromere function may involve modulation of histone H3 levels on chromatin. (A) The decrease of histone H3 levels on chromatin seen in cells over-expressing *rrp1*+ or *rrp2*+ differentially depends on their ATPase and RING domains. Total protein extracts were prepared from cells over-expressing genes encoding wild type and mutated forms of Rrp1-GFP or Rrp2-GFP (B) Data were quantified and are shown as relative intensities of anti-H3, versus anti-GAPDH loading control. Reads were normalised by mean value obtained for vector control samples. (C) Relative intensity of anti-H3 signal, versus Rrp1 or Rrp2 signal, detected with anti-GFP antibodies. Reads were normalised by mean value obtained for *rrp1*+ or *rrp2*+ samples. (D) The decrease of histone H3 levels on chromatin that is induced by DNA damage is partially dependent on the presence of Rrp1 and Rrp2. Total protein extracts were prepared from studied strains incubated in the presence or absence of HU and CPT. (E) Data were quantified and are shown as relative intensities of anti-H3 signal normalised to anti-GAPDH loading controls. Reads were normalised by mean value obtained for the untreated wild type control sample. Western blots were analysed by ImageLab. A minimum of two independent Western blots from three separate protein isolations from respective strains or three different transformants were examined.

It has been reported in yeast that global histone levels are reduced in cells exposed to genotoxic stress (Hauer et al., 2017). We observe similar effect in *S. pombe* wild type strain and demonstrate that it is partially dependent on the presence of Rrp1 and Rrp2 proteins (Fig. 5D,E). These observations are consistent with results obtained for *rrp1*+ or *rrp2*+ over-expression (Fig. 5A,B) and suggest the role of Rrp1 and Rrp2 in regulating nucleosome dynamics.

It has been reported that endogenous Cnp1 (CENP-A) may be redistributed away from centromeric central domain when classical H3 nucleosome assembly is perturbed. Thus, the relative amounts of histones H3 and Cnp1 must be finely balanced in order to ensure proper Cnp1 localisation (Choi et al., 2012). The decrease in histone H3 protein levels (Fig. 6A,B) we observe after prolonged *rrp1*+ and *rrp2*+ overexpression is not accompanied by a statistically significant decrease in Cnp1 levels (Fig. 6A,C), so the ratio of Cnp1 to H3 increases (Fig. 6D). This is especially evident in Rrp1 over-producing cells. Consistently with this, ChIP of endogenously CFP-tagged Cnp1 demonstrates that either *rrp1*+ or *rrp2*+ overexpression leads to an increased Cnp1 enrichment at the core region of centromere (Fig. 6E). We also observed evidence of Cnp1 spreading to neibouring *dg* regions (Fig. 6F). This directly demonstrates that centromere structure is perturbed by *rrp1*+ and *rrp2*+ overexpression. We did not detect in these cells any defect in global regular spacing of nucleosomes using an MNase ladder assay (Fig. S6). This is reminiscent of what was reported for mutants devoid of the CHD1 chromatin remodelers (Pointner et al., 2012; Walfridsson et al., 2007).

**Figure 6.**
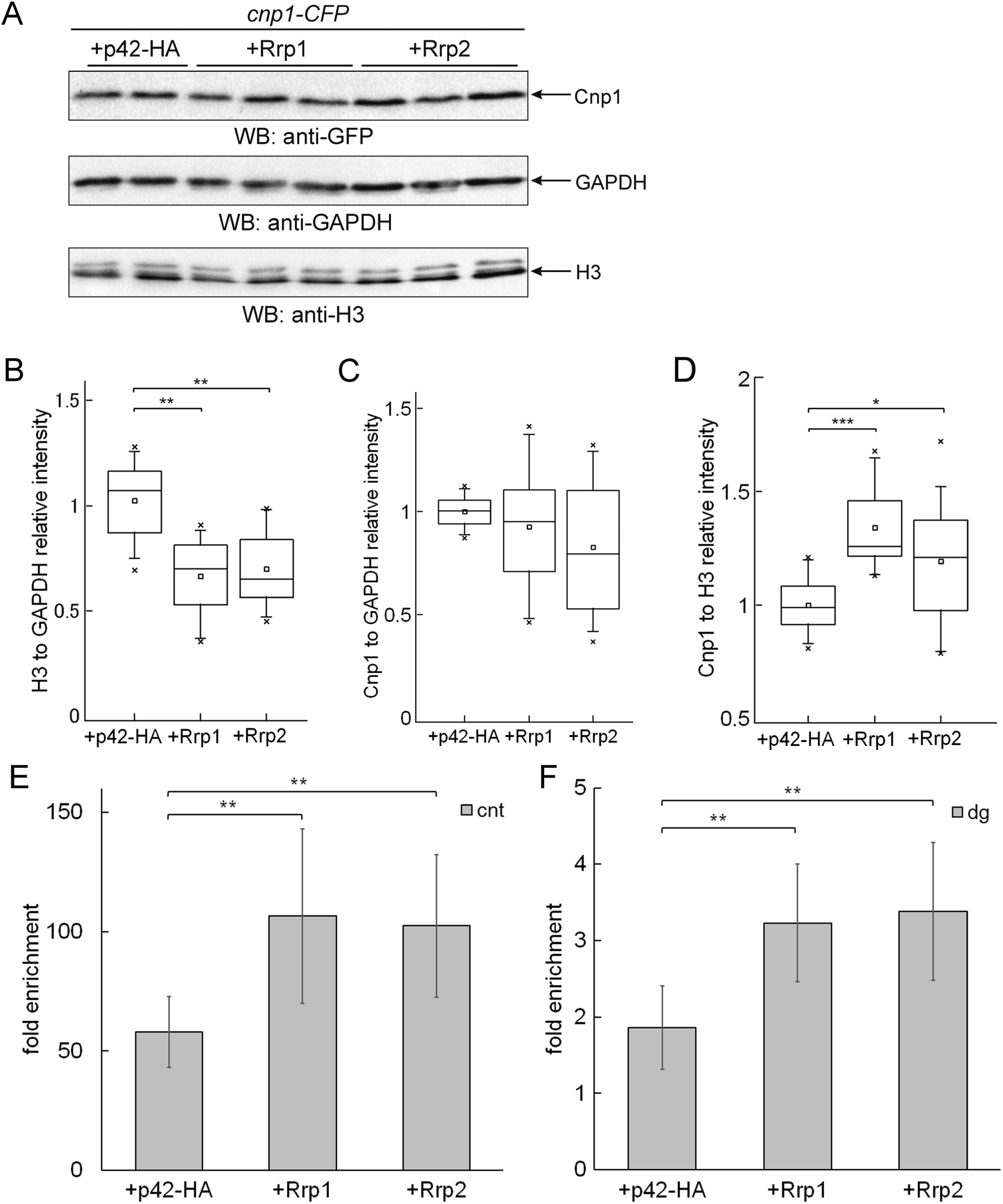
The decrease of histone H3 levels on chromatin caused by over-expression of *rrp1*+ or *rrp2*+ is accompanied by changes in localisation of centromeric histone H3 variant, Cnp1. (A) Total protein extracts were prepared from *cnp1-CFP* strain over-expressing genes for Rrp1-HA or Rrp2-HA. (B) Data were quantified and shown as relative intensities of anti-H3 signal versus anti-GAPDH loading control. (C) Relative intensity of Cnp1 signal, detected with anti-GFP antibodies, versus anti-GAPDH loading control. (D) Relative intensity of Cnp1 signal, versus anti-H3 signal. All reads were normalised by mean value obtained for vector control samples. Western blots were analysed by ImageLab. A minimum of two independent Western blots from three separate protein isolations from three different transformants were examined. (E and F) Endogenous Cnp1-CFP localisation at centromere core (E) and outer repeat (F) regions in cells over-expressing *rrp1*+ or *rrp2*+ was examined by chromatin immunoprecipitation. Fold enrichment is calculated relative to actin control. Primers were located in centromere core (cnt) and outer repeat (dg) regions. Data were obtained from three independent experiments, for each, real time PCR was repeated at least twice. +p42-HA: empty vector control.

Taken together our results suggest that Rrp1 and Rrp2 modulate histone levels on chromatin and that the defects in chromosome segregation and transcriptional silencing observed upon *rrp1*+ and, to some extent, *rrp2*+ overexpression might be due, at least in part, to H3 nucleosome depletion and Cnp1 mislocalisation.

### Rrp2 influences telomere function

*rrp2*+ over-expression generates cell toxicity that is significantly more pronounced than that seen for *rrp1*+ over-expression (Fig. 1A, S4C). However, the growth defect caused by Rrp2 can be uncoupled from transcription silencing effects at the centromere (Fig. 1E). Rrp2 overproduction has recently been shown to result in the accumulation of high-molecular-weight (HMW) SUMO conjugates (Nie et al., 2017) and it has been proposed that the accumulation of Pli1-dependent SUMO chains is toxic (Prudden et al., 2011). In contrast to the overproduction of Rrp2, we observe no detectable increase in HMW SUMO conjugates when Rrp1 was overproduced (Fig. 7A).

**Figure 7.**
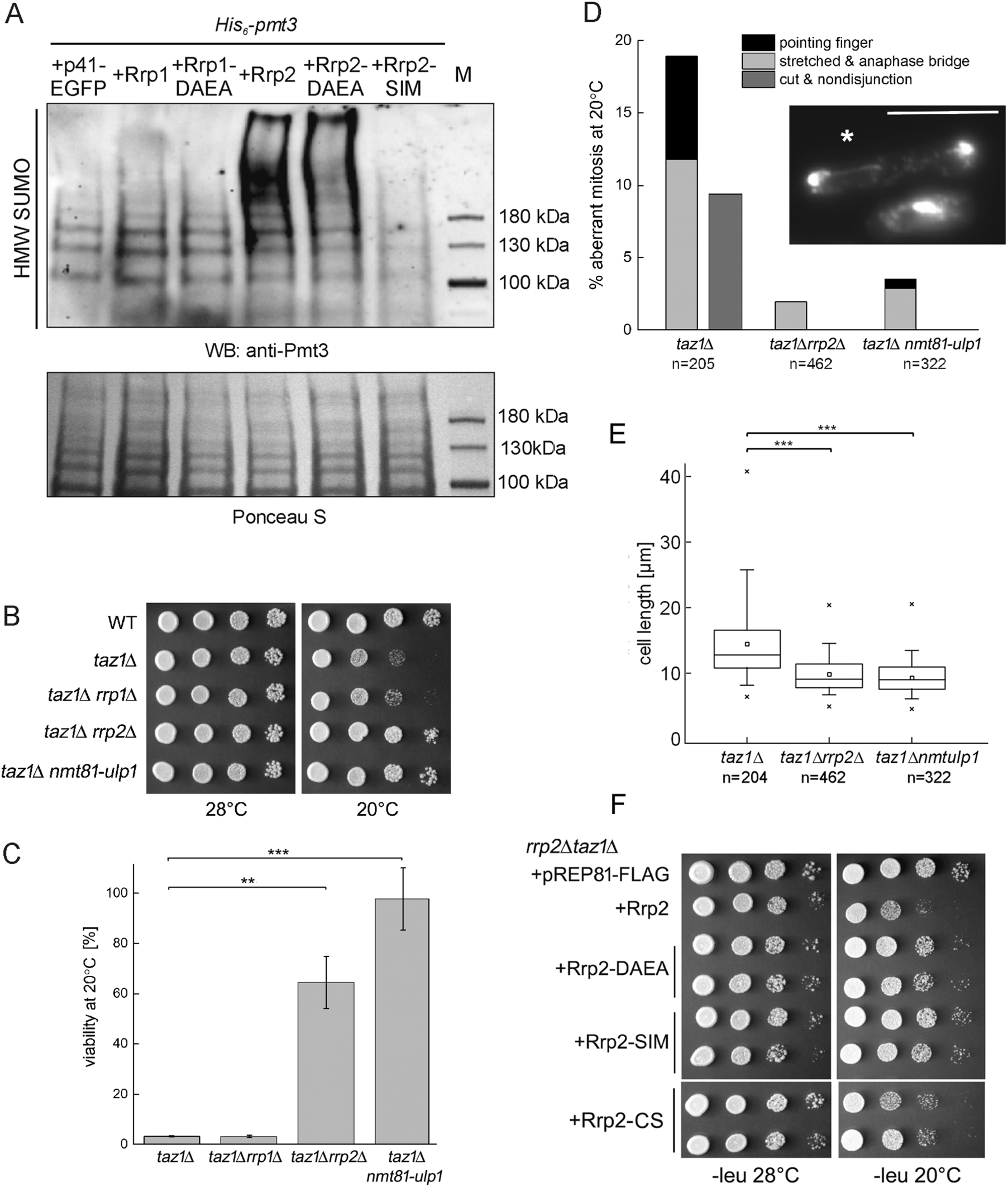
Deletion of *rrp2*+ rescues *taz1*Δ telomere entanglement phenotypes. (A) HMW SUMO conjugates accumulate when *rrp2*+ but not *rrp1*+ is over-expressed and functional Rrp2 SIM motif is essential. The strain with *His*_*6*_-*pmt3* was transformed with empty vector pREP41-EGFP as control and plasmids harbouring genes for wild type or mutated versions of respective proteins. After 24 h growth in minimal medium in absence of thiamine total protein extracts were isolated and subjected to Western blot analysis. SUMOylated proteins were detected with anti-Pmt3 polyclonal serum. Deletion of *rrp2*+ rescues *taz1*Δ mutant’s temperature sensitivity seen as (B) growth inhibition when serial dilutions of tested strains were incubated at 20°C, and (C) the differences in viability of cultures grown at 20°C as compared to 28°C. The error bars represent the standard deviation about the mean values, the experiment was repeated at least three times. (D) Decrease of the characteristic *taz1*Δ mutant anaphase defects (telomere specific pointing finger structure is marked as *) and chromosome breakage and/or missegregation after *rrp2*+ deletion. Cultures of respective strains were grown at 20°C for 2 days and their nuclei stained with DAPI. The experiment was repeated twice and the total number of cells counted for each strain (n) is shown below. Scale bar represents 10 µm. (E) Checkpoint activation, observed as cell elongation, is reduced in the double mutant *taz1*Δ*rrp2*Δ. (F) ATPase activity and SUMO binding are responsible for Rrp2 toxicity in *taz1*Δ mutant. *taz1*Δ*rrp2*Δ double mutant was transformed with empty vector pREP81-FLAG, vector carrying wild type and mutated forms of *rrp2*+ gene and the ability of the constructs to repress growth at 20°C under expression induction conditions was assessed.

The reduction in the levels of HMW SUMO conjugates by overproduction of SUMO protease Ulp1 (Rog et al., 2009) and by deletion of SUMO ligase Pli1, recently shown by (Nie et al., 2017), rescue the cold sensitivity resulting from telomere entanglements generated by aberrant telomere replication in *taz1*Δ cells. As mentioned earlier, the *top2-191* mutation also alleviates *taz1*Δ entanglement phenotypes (Germe et al., 2009). Recently, Rrp2, but not Rrp1, has been demonstrated to antagonize SUMO chain-directed Slx8 ubiquitin ligase activity towards Top2 in a manner dependent on functional Rrp2 ATPase and SIM domains. Loss of Rrp2 was also shown to generate genomic stress in a *top2-191* mutant (Wei et al., 2017). Taken together, these data suggest that Rrp2, but not Rrp1, influences telomere function and that this could explain the differential toxicity observed when over-expressing *rrp2*+ when compared to *rrp1*+.

Consistent with this hypothesis, *taz1*Δ*rrp2*Δ but not *taz1*Δ*rrp1*Δ double mutants grow well at low temperature (Fig. 7B), with viability comparable to that previously reported for over-expression of the Ulp1 SUMO protease in the *taz1*Δ background (*taz1*Δ*nmt81-ulp1*: Fig. 7C). Deletion of *rrp2*+ also reverses the characteristic *taz1*Δ mitotic defects that are a consequence of telomere entanglements: anaphase bridges and “pointing finger” structures, chromosome missegregation events (Fig. 7D) and systemic checkpoint activation resulting in cell elongation are all significantly reduced in the double mutant (Fig. 7E). If the interaction of Rrp2 with Top2 is involved in the role of Rrp2 at telomeres, it would be expected that this toxicity is dependent on the Rrp2 SUMO interaction and translocase activities, but not on functional RING domain (Wei et al., 2017) (see Discussion). Over-expressing *rrp2*+ in the *taz1*Δ*rrp2*Δ mutant, as expected, restored cold sensitivity. In contrast, over-expressing *rrp2-DAE*A and *rrp2-SIM* did not, whereas over-expression of *rrp2-CS* phenocopied *rrp2*+ (Fig. 7F). Thus, as previously shown for the interaction with Top2, Rrp2 SUMO binding and translocase activities are detrimental in cells devoid of Taz1 whereas RING domain is not involved.

*taz1*Δ mutant cells accumulate DNA bridges coated with RPA that cannot be resolved at lower temperatures (Zaaijer et al., 2016). We therefore exploited time lapse microscopy in strains expressing Rad11-GFP to establish if *rrp2*Δ prevented the formation of *taz1*Δ-dependent anaphase bridges, or if such bridges were formed in *taz1*Δ*rrp2*Δ cells, but more efficiently processed. We observed that extensive anaphase bridges arise in both *taz1*Δ and *taz1*Δ*rrp2*Δ cells, but that the number of bridges observed in septated cells is dramatically reduced only in the double mutant (Fig. 8A). These data suggest that in *taz1*Δ mutant cells chromosome bridges persist and lead to chromosome breaks during septum formation (Fig. 8B) and that in *taz1*Δ*rrp2*Δ cells the bridges can be resolved before septation is completed (Fig. 8C). We also show that deletion *rad51*+ rescues the *taz1*Δ mutant’s cold sensitivity (Fig. 8D) and that *taz1*Δ*rad51*Δ double mutant fails to accumulate clusters of Rad11 foci when grown at low temperature (Fig. 8E). This indicates that the formation of telomere entanglements in *taz1*Δ cells results from dysregulation of homologous recombination. Since our data show that deletion of *rrp2*+ is epistatic to *rad51*+ deletion in *taz1*Δ cells (Fig. 8D,E), we propose that Rrp2 is not involved in the formation of *taz1*Δ-dependent telomere entanglements, but inhibits the resolution of these Rad51 dependent structures, which arise illegitimately when telomere ends are destabilized.

**Figure 8.**
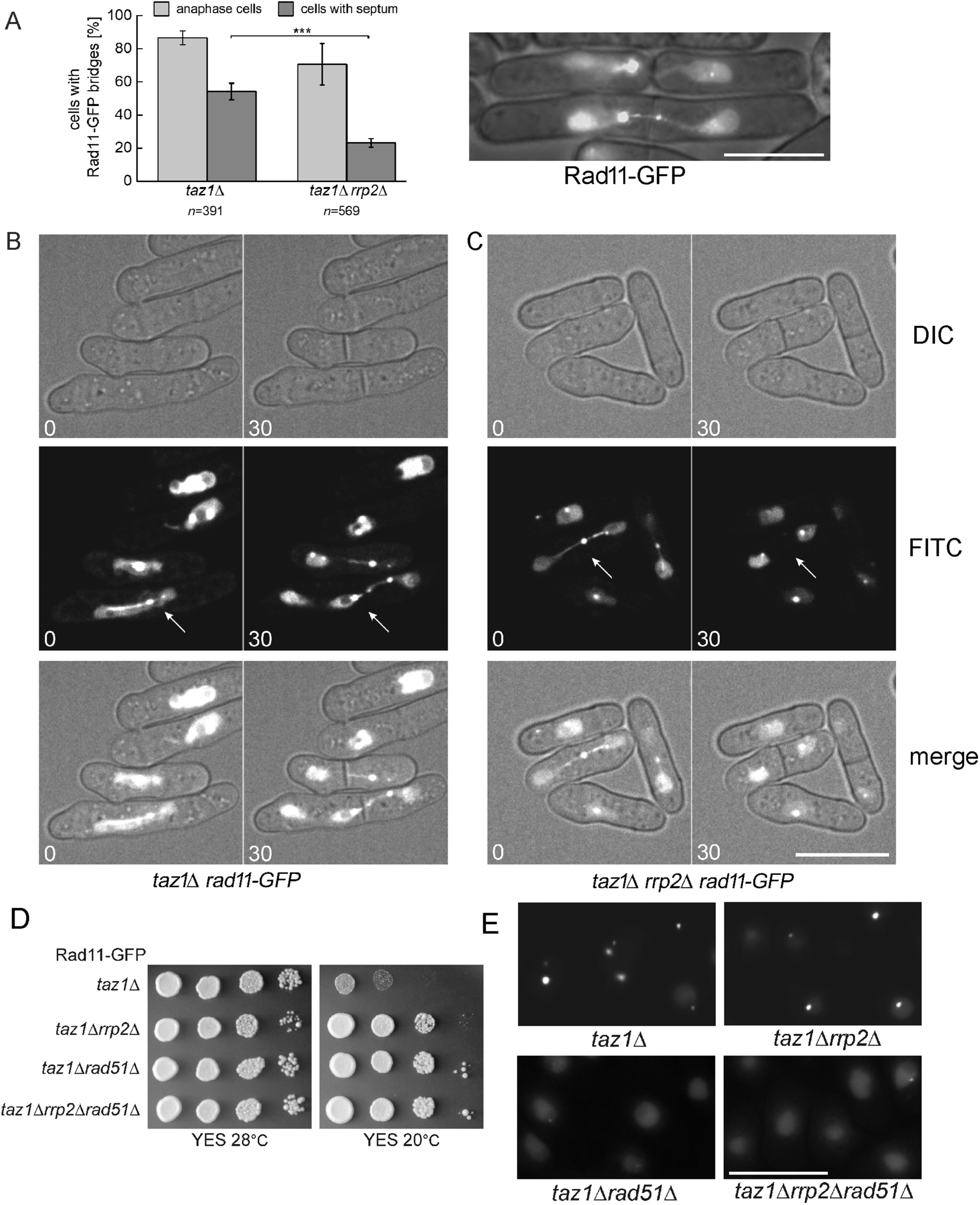
Rrp2 inhibits the resolution of Rad51-dependent telomere entanglements in *taz1*Δ cells. (A) Deletion of *rrp2*+ in the *taz1*Δ mutant reduces the number of chromosome entanglements persisting after septation is completed. To visualize lagging chromosomes *taz1*Δ and *taz1*Δ*rrp2*Δ mutants were analysed in a background of a strain expressing Rad11-GFP form endogenous locus. Cultures were grown at 20°C for 3 days, washed and analysed by fluorescence microscopy. The experiment was repeated four times and the total number of cells counted for each strain (n) is shown. An example of septated cells with unresolved chromosomes is shown at the right. Movie stills showing two different patterns of chromosome segregation during mitosis. White arrows mark irreversible telomere entanglements in (B) *taz1*Δ single mutant, and (C) the ones that can be resolved before septation in a *taz1*Δ*rrp2*Δ double mutant. Images were recorded at the initial time of temperature shift (0) and after 30 minutes (30) of incubation at 20°C. (D) Deletion of *rad51*+ suppresses *taz1*Δ cold sensitivity assayed by growth on plates at 20°C. (E) Chromosome entanglements, assayed by Rad11 foci clusters, are no longer formed in *taz1*Δ*rad51*Δ cells grown at 20°C for 2 days. The experiment was repeated twice and total number of cells counted for each strain was >200. Scale bars represent 10 µm.

We conclude that the improper processing of telomere replication intermediates, which is distinct from the effects of Rrp2 at centromeres, may be the source of the increased Rrp2 overproduction-induced growth defect when compared to Rrp1.

## DISCUSSION

Replication fork progression is often hindered at centromeres and telomeres by secondary structures naturally arising at repetitive sequences. The proteins involved in heterochromatin function include histone modifiers, homologous recombination factors and proteins involved in the replication stress response. More recently, post-translational modifications by ubiquitin and SUMO have also been recognized as important regulators of centromere and telomere integrity (García-Rodríguez et al., 2016; Yalçin et al., 2017). Here we have uncovered roles in centromere and telomere maintenance for two SWI/SNF family members, Rrp1 and Rrp2, that are orthologues of *S. cerevisiae* Uls1, a DNA translocase and STUbL. We showed that over-expression of *rrp1*+ or *rrp2*+ impaired chromosome segregation as evidenced by the formation of mitotic aberrations, increased chromosome instability and viability loss. The viability loss was especially apparent in *rrp2*+ over-expressing cells. Both Rrp1 and Rrp2 formed spontaneous foci in the nucleus and partially co-localized with the Swi6 protein, a heterochromatin marker in fission yeast. Our immunoprecipitation data also indicate that Rrp1 and Rrp2 can bind to centromeric and/or telomeric chromatin. This suggests that the phenotypes conferred by *rrp1*+ or *rrp2*+ over-expression may result from disruption of centromere and/or telomere structure.

Chromosome instability and lagging chromosomes during mitosis can result from aberrant centromere chromatin structure or directly from defective spindle–chromosome interaction (Maruyama et al., 2006). To assess centromere structure and function we therefore examined the silencing of an *ade*+ reporter gene inserted within a *dg* centromeric repeat and assayed for TBZ sensitivity. We found that *rrp1*+ and *rrp2*+ over-expression increased silencing of the *dg*-*ade6*+ gene. What’s more, *rrp1*+ and, to a lesser degree, *rrp2*+ over-expression was able to reverse the de-repression of *dg*-*ade6*+ that is conferred by *swi6*+ deletion. While neither *rrp1*+ or *rrp2*+ deletion or over-expression showed a marked effect on TBZ sensitivity in wildtype cells, *rrp1*+ or *rrp2*+ deletion rescued growth defect of *clr4*Δ mutant, and *rrp1*+ over-expression greatly increased the TBZ sensitivity of *swi6*Δ cells (*rrp2*+ over-expression showed intermediate phenotype). These observations imply that Rrp1 and, to some extent, Rrp2 have roles at the centromere that can influence both its structure and chromosome segregation. This becomes especially evident when centromere structure is disturbed, as in cells lacking proteins Clr4 and Swi6, crucial for heterochromatin formation.

Histone ubiquitylation plays important roles in the DNA damage and replication stress responses as well as in transcriptional silencing (Uckelmann and Sixma, 2017). Thus, the ubiquitin ligase complexes involved are often important for centromere structure maintenance. We showed that the growth defects observed upon *rrp1*+ or *rrp2*+ over-expression are dependent on these protein’s RING domains. We were also able to pull-down ubiquitylated proteins with both Rrp1 and Rrp2 and show that ubiquitylated proteins accumulated in cells over-expressing *rrp1*+ and, to minor extent, *rrp2*+. Taken together, these data suggest that Rrp1 and Rrp2 may indeed have ubiquitin ligase activities.

In *S. pombe* it has previously been shown that histone H2B ubiquitylation, mediated by Rhp6-Brl1/2 complex (RNF20/RNF40 in humans) antagonizes silencing at heterochromatin regions and that in an *H2B-K119R* mutant (that cannot be ubiquitylated) centromeric transcription is strongly repressed (Zofall and Grewal, 2007). We confirmed that the *H2B-K119R* mutation causes transcriptional silencing of *dg-ade6*+ and extended that finding to show that this effect is independent of Swi6 and correlates with a marked increase in TBZ sensitivity. This phenotype is remarkably similar to that which we observed when *rrp1*+ or *rrp2*+ were over-expressed. It has been proposed that, in yeast, loss of H2B ubiquitylation results in nucleosome instability as decreased H2B and H3 levels on chromatin have been observed in H2B-K123R and *rad6*Δ mutants (Chandrasekharan et al., 2009). While we did not detect any effect of *rrp1*+ or *rrp2*+ over-expression on the H2BUbi to H2B ratio, we did find that global histone levels (H2B and H3) are reduced when *rrp1*+ or *rrp2*+ were over-expressed. It is thus possible that a decrease in the total amount of H2BUbi, and/or global histone depletion may influence these common phenotypes.

The depletion of global histone levels, dependent on INO80 nucleosome remodeler, has been recently demonstrated in *S. cerevisiae* to play a role in the response to DNA damage. It was proposed to lead to increased chromosome flexibility and elevated DNA mobility, which enables a more efficient homology search (Hauer et al., 2017). Failure to properly regulate histone levels may be deleterious: increased mobility could enhance access of damaged chromosome to homologous regions on other chromosomes, contributing to DNA translocations and chromosome rearrangements, hallmarks of many cancers. It is thus possible that, through regulation of nucleosome dynamics, Rrp1 and Rrp2 modulate the choice of repair pathway at sites of DNA damage or stalled replication forks. This activity might lead to the Rrp1 and Rrp2 toxicity we observe when their copy number is increased, as well as in specific mutant contexts, as discussed below for Rrp2.

Interestingly, it has been reported that histones H3 and Cnp1 (CENP-A) compete for incorporation into chromatin, such that their relative amounts are important for proper Cnp1 centromeric localisation (Choi et al., 2012). We demonstrate that canonical H3 histone loss in cells over-expressing *rrp1*+ and *rrp2*+ leads to a relative increase in Cnp1 levels, and importantly, to its incorporation in *dg* regions, from which it is normaly excluded. Misincorporation of CENP-A results in transcriptional defects and genomic instability in *S. cerevisiae* (Hildebrand and Biggins, 2016) and leads to defective kinetochore function and chromosome segregation in *S. pombe* (Castillo et al., 2007). Overexpression and mislocalisation of CENP-A can also lead to chromosome instability in human cells (Shrestha et al., 2017) and has been observed in several cancers where it is associated with poor patient survival (Sun et al., 2016; Zhang et al., 2016). The phenotypes of *rrp1*+ and *rrp2*+ over-expression described above: transcriptional repression, problems with DNA segregation leading to TBZ sensitivity and chromosome instability as well a growth defect, could thus at least in part be attributed perturbed centromere structure resulting from mislocalisation of Cnp1.

Chromatin remodelling factors have a well-established role in maintaining appropriate histone levels, ensuring proper Cnp1 localisation to centromeres (Prasad and Ekwall, 2011). Hrp1 (CHD1) and Ino80, facilitate the incorporation of Cnp1 (CENP-A) at core and also at outer repeat regions of the centromere, by actively removing histone H3-containing nucleosomes (Choi et al., 2017; Walfridsson et al., 2007). Fft3 (FUN30) regulates histone H3 levels by modulating higher-order chromatin structures at centromeres (Strålfors et al., 2011). Rrp1 and Rrp2 belong to the Snf2 family of enzymes (Prasad and Ekwall, 2013). We show that for Rrp1 functional ATPase domain is essential for H3 depletion, TBZ sensitivity and silencing, while RING domain is required only for the two latter phenotypes. For Rrp2, both domains are equally important. Both Rrp1 and Rrp2 are STUbL orthologues. *S. cerevisiae* STUbL, Slx5, has been recently found to regulate CENP-A (Cse4) proteolysis (Ohkuni et al., 2016). This raises the possibility that Rrp1 and Rrp2 might be novel chromatin remodelers and/or that they regulate histone dynamics and centromere structure through response to or modulation of protein ubiquitylation. Lending support to this conjecture Rrp2 has been recently identified, together with the histone chaperone Hip1, subunits of the Ino80 complex, components of a Swi2/Snf2 family remodeling complex (Swr1, Swc2), and the nucleosome evictor Fft3 (Fun30) as a factor contributing to recombination hotspot activation during meiosis via a process potentially involving the exchange of individual histone subunits (Storey et al., 2018). Identification of Rrp1 and Rrp2 molecular mode of activity and their putative target(s) remains a challenging task for the future.

*rrp2*+ over-expression leads to greater cell toxicity than that seen when *rrp1*+ is over-expressed. However, TBZ sensitivity of cells over-expressing *rrp2*+ is relatively lower than that induced by *rrp1*+ over-expression. In that respect it is notable that all Rrp2 domains, including the SIM domain, are required for toxicity when *rrp2*+ is over-expressed, whereas SIM motifs are dispensable for the silencing of *dg-ade6*+ reporter gene within the centromere. Also, the increased loss of viability caused by *rrp2*+ over-expression is dependent on the presence of Rad51 recombinase. Thus, the growth defect induced by *rrp2*+ over-expression can be partially separated from its role in centromere structure, demonstrating that Rrp2 has other function(s) that are important for viability loss when *rrp2*+ is over-expressed. Our data further show that these are dependent on the ATPase and SIM domains and function independently of Rrp1. In accord with these observations it has recently been shown that Rrp2, but not Rrp1, protects cells from Top2-induced DNA damage (Wei et al., 2017).

Based on the above, and on the known function of Top2 in telomere replication, we hypothesized that Rrp2, but not Rrp1, is involved in telomere maintenance and that this could explain the differential toxicity of *rrp1*+ and *rrp2*+ over-expression. In support of this, we showed that deletion of *rrp2*+, but not of *rrp1*+, reversed *taz1*Δ mitotic defects. Using time lapse microscopy we demonstrated that telomere entanglements arise both in *taz1*Δ and in *taz1*Δ*rrp2*Δ cells but that, in the presence of Rrp2, their resolution is prevented. We also showed that Rrp2 SUMO binding and translocase activities were necessary for Rrp2 toxicity in the *taz1*Δ mutant background. We thus propose that the interaction with Top2 described in (Wei et al., 2017) underpins the role of Rrp2 at the telomeres: in *rrp2*Δ mutants Top2 is not protected from STUbL degradation and this increases the probability of exposing a DSB if Top2 is degraded while still at the Top2cc stage (Wei et al., 2017). Similarly, *top2-191* mutant (Germe et al., 2009), which is trapped longer at the Top2cc stage during catalytic cycle, faces higher risk of being inadvertently degraded by STUbL, thus exposing DNA breaks. While these outcomes are generally avoided in otherwise wild type cells, in *taz1*Δ (where telomere separation during anaphase is hindered by entanglement) introducing breaks into telomeric DNA likely allows the separation of the chromosomes. This would prevent chromosome arm breakage due to incomplete DNA segregation and subsequent septation in *taz1*Δ mutant and thus lead to the rescue of the *taz1*Δ growth defect seen in both *top2-191* (Germe et al., 2009) and *rrp2*Δ backgrounds.

Taken together our data suggest that, even though preventing untimely removal of Top2cc complexes from DNA by Rrp2 ensures genetic stability in wildtype cells and protects from Top2 poisons (Wei et al., 2017), it can be detrimental in specific mutant contexts (such as in *taz1*Δ) and when exacerbated in wildtype cells by protein overproduction. We show that telomere entanglements do not form in *taz1*Δ*rad51*Δ cells which is consistent with lower toxicity of *rrp2*+ overexpression seen in *rad51*Δ mutant. Interestingly, overproduction of human SLX4 has also been shown to be toxic in cells exposed to global replication stress and this effect was dependent on the protein’s SUMO ligase activity. However, the same ligase activity was essential for the resolution of mitotic interlinks at chromosome fragile sites (Guervilly et al., 2015). This is another example consistent with a model whereby diverse genomic regions are differentially regulated, with the controlled promotion of DSB formation crucial in difficult-to-replicate regions of the genome.

Accumulating data clearly demonstrate that DNA metabolic processes in vulnerable regions of the genome must be finely tuned to ensure their stability. The activities involved are often different to those invoked in response to global DNA damage or replication stress. Consistent with this, Rrp2 has been shown to activate a specific meiotic recombination hotspot without affecting basal recombination levels (Storey et al., 2018). In *S. cerevisiae* it has been demonstrated that the relocation of DSBs flanked by repeated sequences (such as those found in telomeres) to nuclear pores depends on Uls1, an Rrp2 orthologue, but not on general stress response STUbL, Slx5 (Marcomini et al., 2018). Interestingly, the opposite is true for DSB appearing throughout the genome (Horigome et al., 2016).

The consequences of deleting genes encoding proteins that modulate DNA metabolic processes at specific genomic regions can often go undetected in experimental systems because general DNA damage responses can substitute at the expense of modest defects in genome stability. However, dysregulation of these gene’s activity through their over-expression, such as described here for Rrp1 and Rrp2, is likely to perturb the system more dramatically and generate abnormal replication and/or repair intermediates, leading to more extensive genomic instability and a consequent viability loss. It is of note that also in humans, even if mutations in some genes, like *RECQ5* for example, are not directly associated with predisposition to cancer or genetic disease, their amplification and increased expression is found in many tumors, and has been shown to redirect repair pathways and lead to genomic instability (Olson et al., 2018).

## MATERIALS AND METHODS

### Yeast strains, plasmids and general methods

Strains and plasmids used in this study are listed in Tables S1 and S2, respectively. Media used for *S. pombe* growth were as described (Moreno et al., 1991). Yeast cells were cultured at 28°C in complete yeast extract plus supplements (YES) medium or glutamate-supplemented Edinburgh minimal medium (EMM). Thiamine was added where required (5 μg/mL), as were geneticin (ICN Biomedicals) (100 µg/mL), nurseotricin (Werner Bioagents) (200 μg/mL) and hygromycin (Sigma-Aldrich). For YES low ade plates, the concentration of adenine was reduced 10-fold. pREP81-FLAG vector and plasmids carrying wild type and mutated forms of *rrp1*^+^ and *rrp2*^+^ were constructed using Gibson Assembly Cloning Method/ Gibson Assembly® Cloning Kit (NEB). All primers used to amplify gene sequences by PCR are listed in Table S3. Amplified fragments were cloned into NdeI and BamHI digested pREP81 vector. After Gibson cloning inserts were cut by NdeI and SmaI digestion and cloned into pREP41-EGFP plasmid. Plasmid over-expressing *rrp2-SIM* was obtained by cloning a *rrp2*+ coding sequence from pDUAL-Prrp2-GFP-Rrp2-SIM(1-6)*, a gift from Li Lin Du (Wei et al., 2017) into pREP41-EGFP.

### Whole protein extract analysis

Protein extracts were prepared by the trichloroacetic acid (TCA) method. Briefly, after 24 hour of induction of *nmt* promoter by removal of thiamine from media, mid-logarithmic cells (∼10^8^) of indicated strains were harvested and lysed with lysis buffer (2 M NaOH, 7% β-mercaptoethanol). Total protein was precipitated by adding 50% TCA. Pellet was then resuspended in 1 M Tris at pH 8 and 4x Laemmli buffer was added (250 mM Tris-HCl, pH 6.8, 8% SDS, 20% glycerol, 0.02% Bromophenol blue, 7% β-mercaptoethanol). Obtained samples were analysed by SDS-PAGE and Western blotting using anti-GFP (Roche, 11814460001), anti-FLAG (Sigma-Aldrich, F1804), anti-H3 (Abcam, ab1791) or anti-GAPDH (loading control, Invitrogen, MA5-15738) antibodies. Blotted membranes were stained with the Ponceau S (Sigma-Aldrich) to detect total proteins. For protein quantification Image Lab (Western blots) or ImageJ software (Ponceau S staining) was used. Relative intensity was calculated by dividing sample intensities by the mean of control intensities obtained for each blot (details for each experiment are provided in figure captions). For each experiment data from two different transformants from two independent protein isolations were analysed.

### Detection of high-molecular weight SUMO conjugates

Protein extracts for identifying high-molecular weight SUMO-conjugates were prepared according to (Nie et al., 2017) with following modifications. After 24 hour induction of *nmt* promoter by removal of thiamine from media, mid-log cells (∼2×10^8^) were washed with STOP buffer (10 mM EDTA, 50 mM NaF, 150 mM NaCl) and pellets were frozen in liquid nitrogen. Cells were resuspended in 200 μL of 20% TCA with 200 μL of glass beads (Roth) and subsequently disrupted by bead beating. Next, 400 μL of 5% TCA was added, lysate was separated from the beads and centrifuged at 16,000 *g* for 5 min at 4°C. The pellet was washed twice with 0.1% TCA. The precipitated proteins were resuspended in 8 M urea, 50 mM Tris, pH 8.5, 150 mM NaCl. After estimation of protein concentration by measurement of absorbance at 280 nm, 2x loading buffer was added (6 M urea, 62.5 mM Tris-HCl pH 6.8, 2% SDS, 20% glycerol, 0.01% bromophenol blue, 3.5% β-mercaptoethanol) and samples were analysed by SDS-PAGE using 4-20% gradient Mini Protean TGX Precast Gel (Bio-Rad). Membrane was visualized for total protein with Ponceau S (Sigma-Aldrich) and Western blot was performed using anti-Pmt3 polyclonal serum (gift from Felicity Watts).

### Spot assays

Cells were grown to mid-log phase, then serially diluted by 10-fold and 2 µL aliquots were spotted onto relevant plates (YES or EMM) without drug or plates containing thiabendazole (TBZ) at concentrations indicated at each figure. Plates were incubated for 3-5 days in 28°C (unless stated otherwise) and photographed. All assays were repeated at least twice. TBZ was added to the plates at the concentrations.

### Survival assay

Cells were grown for 48 hours in minimal medium with (repressed conditions) or without thiamine (over-expression) at 28°C. 500 µL aliquots were collected, serially diluted and plated onto YES plates to determine the number of viable cells. Plates were incubated for 3-5 days at 28°C. The viable cells were counted and percentage of survival was calculated. For temperature survival, cells were grown to mid-log phase in rich medium in 28°C or 20°C. Samples were collected and diluted in the same way as above, plated onto YES plates and incubated for 3-5 days at 28°C. The percentage of survival in 20°C was calculated against the 28°C control.

### Chromosome loss

Indicated strains where streaked to single colonies on EMM low Ade plates (adenine concentration reduced to 7.5 mg/L) with thiamine and a single white colony was inoculated in EMM without thiamine and incubated for 48 h at 28°C. Then cultures were appropriately diluted, plated on YES low Ade plates and incubated for 3-4 days at 28°C. Percentage of red to white colonies was then calculated.

### Fluorescence microscopy

To determine the foci formation of Rrp1 and Rrp2 proteins, their co-localisation with Swi6 protein and Rad11, and influence of *rrp1*+ or *rrp2*+ over-expression on Rad11-GFP localization appropriate strains were grown for 24 h in EMM medium without thiamine. 1 mL of culture was harvested, washed with water and subjected to fluorescent microscopy analysis. For Rrp1 and Rrp2 foci images were captured under 100x magnification using Axio Imager A.2 (Carl Zeiss) with Canon digital camera, and analysed with Axiovision rel. 4.8. For co-localisation experiments, data were collected under 63x magnification with confocal microscope Leica TCS SP8 (Leica Microsystems) equipped with Leica HyD SP detector, and analysed with LAS X 3.3.0.

For examination of mitotic defects induced by *rrp1*^+^ or *rrp2*^+^ over-expression samples taken from cultures grown for 48 hours in EMM medium without thiamine. were washed and fixed in 70% ethanol. After rehydration, cells were stained with 1 mg/mL 4’,6-diamidino-2-phenylindole (DAPI) and 1 mg/mL p-phenylenediamine in 50% glycerol and examined by fluorescence microscopy with Axio Imager A.2 (Carl Zeiss).

To observe Rad11-GFP foci in *taz1*Δ and *rrp2*Δ*taz1*Δ both mutants were grown to mid-logarithmic phase in EMM, 20°C. Cells were then centrifuged and resuspended in 1 mL of fresh EMM. A drop of 1 µL was spotted on the layer of 1.4% agarose in filtered EMM covering a Thermo Scientific slide (ER-201B-CE24). Z-stack pictures were captured using a 3D microscope (LEICA DMRXA) equipped with a CoolSNAP monochromic camera (Roper Scientific) under 100X magnification, exposure time for GFP 500 ms, with METAMORPH software. Image analysis was performed with ImageJ software.

For time-lapse movies, cells were processed in the same way as for the snapshot microscopy and visualized with a Nikon inverted microscope equipped with the Perfect Focus System, a 100X/1.45-NA PlanApo oil immersion objective, Yokogawa CSUX1 confocal unit, Photometrics Evolve512 EM-CCD camera and a laser bench (Errol) with 491 nm diode laser, 100 mX (Cobolt). Images were captured every 15 s with 500 ms exposure time for GFP at 15% of laser power using METAMORPH software. Movies were mounted and analysed using ImageJ software. Image acquisition with a LEICA DMRXA 3D microscope and Nikon inverted microscope were performed on the PICT-IBiSA Orsay Imaging facility of the Institut Curie.

### Histone loss upon DNA damage treatment

Cultures of wild type and mutant cells were grown in YES at 28°C to OD 0.4-0.7 and split into two tubes. 12 mM HU and 20 µM CPT was added to one tube (the other serving as an untreated control) and incubation continued at 28°C for 4 hours. Total protein was then isolated and subjected to Western blot analysis.

### Chromatin immunoprecipitation (ChIP)

Experiments were performed as described in (Ait Saada et al., 2017) with small modifications. Briefly, yeast cultures were grown to logarithmic phase in YES medium or for 24 hours in EMM medium without thiamine and 10^9^ cells were pelleted and resuspended in PBS with 2.5 mg/ml DMA with 0.25% DMSO and incubated 45 min with shaking at RT. Cells were pelleted, washed with PBS and incubated with 1% formaldehyde for another 15 min. Glycine was added to neutralize formaldehyde. Cells were then pelleted, washed with PBS, frozen in liquid nitrogen and stored at −80°C. After disrupting cells by bead beating in ChIP lysis buffer (50 mM HEPES pH 7.4; 140 mM NaCl; 1 % Triton X100; 0.1% Na-deoxycholate) with PMSF (1 mM) and protease inhibitor (Complete EDTA –free protease inhibitor cocktail, Roche) 10 cycles of sonication were performed: 20 seconds ON and 60 seconds on ice using water ultrasonicator (LABART). Before immunoprecipitation, input samples were taken as a control, then anti-GFP antibody (Life Technologies, A-11122) was added to each sample. Samples were incubated for 1 h at 4°C, after that 20 μL of Dynabeads Protein G (Thermo Fisher) was added per sample and incubated over night at 4°C. After washing steps and de-crosslink for 2 h at 65°C DNA was recovered by ethanol precipitation. Purified DNA was subjected to further analyses by qPCR using StepOne (Thermo Fisher) thermocycler. Target and control (actin) primers used are listed in Table S3.

Data were collected from at least one qPCR performed on DNA from four independent biological experiments. The Ct value (number of cycles required for the fluorescent signal to cross the threshold) from input samples (Ct_(IN)_) and ChIP samples (Ct_(ChIP)_) was recorded by StepOne Software. Following formula was used to calculate percent enrichment of the amount of protein binding to a target locus over actin: (100×(1/2^(Ct_(ChIP)_-Ct_(IN)_))_target_/(100×(1/2^(Ct_(ChIP)_-Ct_(IN)_))_actin_.

### Immunoprecipitation

Yeast cultures were incubated for 24 h to mid-log phase in EMM medium without leucine and thiamine. Cells (∼3×10^9^) were centrifuged and frozen in liquid nitrogen. Pellets were resuspended in 300 μL of lysis buffer (50 mM HEPES-KOH pH 7, 50 mM KOAc, 5 mM MgOAc, 0.1% NP-40, 10% Glycerol, 1 mM DTT, 0.5 mM PMSF, cOmplete(tm) EDTA-free Protease Inhibitor Cocktail (Roche), 20 mM β-glycerolphosphate) and disrupted by bead beating. Subsequently lysate was centrifuged at 20,000g for 10 min at 4°C. Supernatant was incubated with 30 μL of FLAG M2 agarose (Sigma-Aldrich, A2220) for 3 h at 4°C. After washing three times with lysis buffer, bound proteins were eluted using 100 μL of FLAG peptide (100 μg/mL, Sigma-Aldrich, F4799), and then with 100 μL of 2x Laemli buffer (62.5 mM Tris-HCl, pH 6.8, 2% SDS, 20% glycerol, 0.01% bromophenol blue, 3.5% β-mercaptoethanol). Both eluates were then combined and analyzed by SDS-PAGE and Western blotting using anti-Ub antibody (Abcam, [P4D1] ab95530).

### His-Ub pull down

Yeast cultures were incubated for 24 h in EMM without leucine and thiamine to induce expression of His-tagged ubiquitin and Rrp1 or Rrp2 and protein was isolated as described (Khmelinskii et al., 2014) with modifications. Mid-log cells (∼10^9^) were harvested, washed with 20% TCA and then frozen in liquid nitrogen. Pellets were resuspended in 300 µL of 20% TCA and disrupted by bead beating. Obtained cell lysates were centrifuged for 10 min. at 14,000 rpm, 4°C. TCA was removed and pellets were dissolved in 1 mL of purification buffer (6 M guanidium-Cl, 100 mM Tris pH 9, 300 mM NaCl, 0.2% Triton X-100, 5 mM chloroacetamide, 10 mM imidazole) and 40 µL of TALON beads (Clontech) was added. After rotating for 90 min at RT, beads were washed twice with wash buffer I (8 M urea, 100 mM Na-phosphate buffer pH 7, 300 mM NaCl, 0.2% Triton X-100, 5 mM chloroacetamide, 5 mM imidazole) and twice with wash buffer II (8 M urea, 100 mM Na-phosphate buffer pH 7, 300 mM NaCl, 0.2% SDS, 0.2% Triton X-100, 5 mM chloroacetamide, 5 mM imidazole). During the last wash beads were transferred into a Spin-X tube (Costar, cellulose acetate, 0.45μm), centrifuged for 2 min at 5000 rpm and eluted with elution buffer (8 M urea, 100 mM Na-phosphate buffer pH 6.5, 300 mM NaCl, 0.2% SDS, 0.2% Triton X-100, 5 mM chloroacetamide, 250 mM imidazole). For WB analysis 4x Laemli buffer (250 mM Tris-HCl, pH 6.8, 8% SDS, 20% glycerol, 0.02% bromophenol blue, 7% β-mercaptoethanol) was added, samples were incubated for 20 min at 30°C and subsequently analyzed by SDS-PAGE and Western blotting using anti-Ub antibodies (Abcam, [P4D1] ab95530).

### MNase digestion

The MNase ladder assay was performed according to (Lantermann et al., 2009). After 24 hour induction of *nmt* promoter by removal of thiamine, mid-log cells were crosslinked with 0.5% formaldehyde for 20 minutes. Cell wall was removed by digestion with Zymolase T100 in S buffer (50 mM Tris-HCl pH 7.4, 1 M sorbitol, 10 mM 2-mercaptoethanol) for 1 h. Spheroplasts were resuspended in NP buffer (1 M sorbitol, 50 mM Tris-HCl pH 7.4, 5 mM MgCl_2_, 1 mM CaCl_2_, 0.75% NP-40) and divided into 100 μl samples. Microccocal nuclease (Sigma-Aldrich) was added to samples to the final concentration of 5 U/ml and, after incubation for indicated times, the reaction was stopped by the addition of buffer containing 0.35 M EDTA, 3% SDS, and 1.5 mg/ml proteinase K. Digested DNA was isolated using phenol:chloroform:isoamyl alcohol, washed with ethanol, gently resuspended in MQ water and treated with 0.01 mg/ml RNAseA (Sigma-Aldrich) for 30 minutes in 37°C. Samples were run on 1.5% agarose gel in TAE buffer. Bands were visualized by SimplySafe staining.

### Statistical data analysis

In all box and whiskers plots boxes represent the range from 25 to 75%, whiskers— the range from 5 to 95%, lines dividing the boxes—the median and full squares—the mean value. The error bars represent the standard deviation about the mean values. Student’s t-test was used to calculate the P-values (* 0.01 < P-value ≤ 0.05, ** 0.001 < P-value ≤ 0.01, *** P-value ≤ 0.001).

## ACKNOWLEDGMENTS

We thank Robin Allshire, Julie Cooper, Jason Tanny and C. David Allis, Jo Murray, Susan Forsburg and Benoit Arcangioli for providing strains, Hiroshi Iwasaki for strains and Ub plasmid, Nick Boddy and Minghua Nie for strains and sharing the protocol for isolation of SUMOylated proteins, Felicity Watts for the kind gift of strains and anti-Pmt3 serum and Wojciech Bialek for reagents. We thank Natalia Trempolec and Joanna Morcinek-Orłowska for involvement in the initial stages of this project. We are grateful to Li Lin Du for sharing unpublished data and providing Rrp2 and Rrp2-SIM plasmids, and to Sarah Lambert for comments on the manuscript and hosting Kamila for a research project in her laboratory.

