## Supplementary materials for "*S. pombe* DNA translocases Rrp1 and Rrp2 have distinct roles at centromeres and telomeres that ensure genome stability"

**Table S1** Strains used in this study

| Strain | Genotype | Reference |
|---|---|---|
| YA254 (WT) | ura4-D18, leu1-32, his3-D1, arg3-D1, h90 | (Akamatsu et al., 2003) |
| *rrp1*Δ-1 | rrp1D::kanMX6, ura4-D18, leu1-32, his3-D1, arg3-D1, h90 | lab stock |
| *rrp2*Δ-1 | rrp2D::kanMX6, ura4-D18, leu1-32, his3-D1, arg3-D1, h90 | lab stock |
| *rrp1*Δ | rrp1D::natMX, his3-D1, leu1-32, ura4-D18, arg3-D1, h90 | lab stock |
| *rrp2*Δ | rrp2D::natMX, his3-D1, leu1-32, ura4-D18, arg3-D1, h90 | lab stock |
| *rrp1Δ rrp2*Δ | rrp1D::kanMX6, rrp2D::natMX, his3-D1, leu1-32, ura4-D18, arg3-D1, h90 | lab stock |
| BX5 | $His_6$-pmt3, ura4-D18, leu1-32, ade6-210, h90 | (Xhemalce et al., 2004) |
| AMC377 | rad11-GFP:: kanMX6, ura4-D18, leu1-32, his3-D1, h+ | lab stock |
| GFP-Rrp1 | loxP-GFP-Rrp1-loxM, ura4-D18, leu1-32, h+ | a |
| Rrp2-GFP | Rrp2-GFP::loxP kanMX6 loxM, ura4-D18, leu1-32, ade6-704, h+ | lab stock |
| CJ01 | ura4-D18, leu1-32, his3-D1, arg3-D1, ade6-m210, Chr16 ade6-m216, h+ | b |
| *clr4*Δ | clr4::LEU2, ura4-D18, ade6-210, h+ | c |
| *clr4*Δ-1 | clr4::LEU2, his3-D1, ura4-D18, arg3-D1, leu1-32, h- | a |
| *clr4Δrrp1*Δ | rrp1:: natMX, clr4::LEU2, his3-D1, ura4-D18, arg3-D1, leu1-32, swi+/- | a |
| *clr4Δrrp2*Δ | rrp2:: natMX, clr4::LEU2, his3-D1, ura4-D18, arg3-D1, leu1-32, swi+/- | a |
| YA728 | swi6-ECFP, his3-D1, ura4-D18, arg3-D1, leu1-32, Msmt-0 | d |
| FY3138 | ade6-DN/N, leu1-32, ura4-D18, otr1R(SphI)::ade6+, h+ | (Ekwall et al., 1999) |
| FY3138 *rrp1*Δ | rrp1D::kanMX6, ade6-DN/N, leu1-32, ura4-D18, otr1R(SphI)::ade6+, h+ | a |
| FY3138 *rrp2*Δ | rrp2D::kanMX6, ade6-DN/N, leu1-32, ura4-D18, otr1R(SphI)::ade6+, h+ | a |
| FY3138 *swi6*Δ | swi6D::natMX, ade6-DN/N, leu1-32, ura4-D18, otr1R(SphI)::ade6+, h+ | a |
| FY3138 *swi6Δ rrp1*Δ | swi6D::natMX, rrp1D::kanMX6, ade6-DN/N, leu1-32, ura4-D18, otr1R(SphI)::ade6+, h+ | a |
| FY3138 *swi6Δ rrp2*Δ | swi6D::natMX, rrp1D::kanMX6, ade6-DN/N, leu1-32, ura4-D18, otr1R(SphI)::ade6+, h+ | a |
| FY3138 *clr4*Δ | clr4::LEU2, ade6-DN/N, leu1-32, ura4-D18, otr1R(SphI)::ade6+, h+ | a |
| *rad57*Δ | rad57D::his3+, ura4-D18, leu1-32, his3-D1, arg3-D1, Msmt0 | (Akamatsu et al., 2003) |
| *sfr1*Δ | sfr1D::arg3, arg3-D4, ura4-D18, leu1-32, h- | lab stock |
| *swi5*Δ | swi5D::his3+, ura4-D18, leu1-32, his3-D1, arg3-D1, swi+/- | (Akamatsu et al., 2003) |
| *rrp1Δ rad57*Δ | rrp1D::kanMX6, rad57D::his3+, ura4-D18, leu1-32, his3-D1, arg3-D1, Msmt0 | lab stock |

| | | |
|---|---|---|
| *rrp2*Δ *rad57*Δ | rrp1D::kanMX6, rad57D::his3+, ura4-D18, leu1-32, his3-D1, arg3-D1, Msmt0 | lab stock |
| JTB229 | htb1-K119R-FLAG::kanMX6, ade6, h+ | (Tanny et al., 2007) |
| JTB229-1 | htb1-K119R-FLAG::kanMX6, leu1-32, ade6, his3-D1, h90 | a |
| JTB229 *swi6*Δ | htb1-K119R-FLAG::kanMX6, swi6D:natMX, otr1R(SphI)::ade6+, leu1-32, ura4-D18, swi+/- | a |
| JTB62 | htb1-FLAG::kanMX6, ade6, h+ | (Tanny et al., 2007) |
| JTB62-1 | htb1-FLAG::kanMX6, leu1-32, ura4-D18, ade6, his3-D1, h- | a |
| JTB62 *swi6*Δ | htb1-FLAG::kanMX6, swi6D::natMX, otr1R(SphI)::ade6+, leu1-32, ura4-D18, swi+/- | a |
| JTB62 *rrp1*Δ | htb1-FLAG::kanMX6, rrp1::natMX, leu1-32, ura4-D18, ade6, his3-D1, h90 | a |
| JTB62 *rrp2*Δ | htb1-FLAG::kanMX6, rrp2::natMX, leu1-32, ade6, his3-D1, h90 | a |
| FY 4229 | CFP-cnp1::kanMX6, ura4-D18, rad22::YFP-kanMX6, sad1::sRed-Leu2+, h? | (Li et al., 2013) |
| KK821 | CFP-cnp1::kanMX6, ade6-704, ura4-D18, leu1-32, h? | a |
| JCF1770 | taz1::hygMX, pli1::kanMX6, ura4-D18, his3-D1, leu1-32, ade6-210, h- | (Rog et al., 2009) |
| JCF1779 | taz1::ura4+, nmt81-ulp1::kanMX6, ura4-D18, his3-D1, leu1-32, ade6-210, h- | (Rog et al., 2009) |
| *taz1*Δ | taz1::ura4+, his3-D1, leu1-32, ura4-D18, arg3-D1, h- | a |
| *taz1*Δ *rrp1*Δ | taz1::ura4+, rrp1::natMX, his3-D1, leu1-32, ura4-D18, arg3-D1, h90 | a |
| *taz1*Δ *rrp2*Δ | taz1::ura4+, rrp2::natMX, his3-D1, leu1-32, ura4-D18, arg3-D1, h90 | a |
| *taz1*Δ Rad11-GFP | rad11-GFP::kanMX6, taz1::ura4+, his3-D1, leu1-32, ura4-D18, ade6-210, h+ | a |
| *taz1*Δ *rrp2*Δ Rad11-GFP | rad11-GFP::kanMX6, taz1::ura4+, rrp2::natMX, his3-D1, leu1-32, ura4-D18, h+ | a |

a - this work
b - a gift from Jo Murray
c - a gift from Felicity Watts
d - a gift from Hiroshi Iwasaki

**Table S2** Plasmids used in this study

| Plasmid | Reference |
|---|---|
| pREP41-EGFP | (Craven et al., 1998) |
| pREP41-EGFP-Rrp1 | a |
| pREP41-EGFP-Rrp1-FLAG | a |
| pREP41-EGFP-Rrp1-DAEA-FLAG | a |
| pREP41-EGFP-Rrp1-CS-FLAG | a |
| pREP41-EGFP-Rrp2 | a |
| pREP41-EGFP-Rrp2-FLAG | a |
| pREP41-EGFP-Rrp2-DAEA-FLAG | a |
| pREP41-EGFP-Rrp2-SIM-FLAG | a |
| pREP41-EGFP-Rrp2-CS-FLAG | a |
| pREP42-EGFP | b |
| pREP42-EGFP-Rrp1 | b |
| pREP42-EGFP-Rrp2 | b |
| pREP41-mCherry | b |
| pREP41-mCherry -Rrp1 | b |
| pREP41-mCherry -Rrp2 | b |
| pREP42-HA | (Craven et al., 1998) |
| pREP42-HA-Rrp1 | b |
| pREP42-HA-Rrp2 | b |
| pREP81-FLAG | b |
| pREP81-Rrp1-FLAG | a |
| pREP81-Rrp2-FLAG | a |
| pREP81-EGFP-Rrp2 | b |
| pDUAL-Prrp2-GFP-Rrp2-SIM(1-6)* | (Wei et al., 2017) |
| pYK788 pREP1-His-Ubi1 | c |

a - this study
b - laboratory stock
c - a gift from Hiroshi Iwasaki

**Table S3** Primers used in this study:

| Cloned gene | Primer name | Primer sequence |
|---|---|---|
| FLAG | REP3flag_a_r | ctttatcatcgtcgtccttgtagtcggatcctctagagtcgacatatgatttaac |
| | REP3flag_b_f | tacaaggacgacgatgataaagactacaaggacgacgatgataaagacta |
| | REP3flag_c_r | gggtcatttatcatcgtcgtccttgtagtctttatcatcgtcgtccttgtagt |
| | REP3flag_d_f | caaggacgacgatgataaatgacccgggtaaaaggaatgtctcccttgccagtac |
| Rrp1-FLAG | Rrp1_fwd | caactaattattcgaaacggaattcgaaacgATGGATTCATTGTCTGCATATC |
| | Rrp1_rev | tttaaatggccggccggtaccTCATGAATTAAGCCCAAATAG |
| Rrp1-DAEA-FLAG (D397A; E398A) | Rrp1_fwd | caactaattattcgaaacggaattcgaaacgATGGATTCATTGTCTGCATATC |
| | Rrp1D397A_rev | tatgtgcggcCGCTAGAACAATGCGATAC |
| | Rrp1E398A_fwd | tgttctagcgGCCGCACATACCATTCGT |
| | Rrp1E398A_rev | tttaaatggccggccggtaccTCATGAATTAAGCCCAAATAGATATAG |
| Rrp1-CS-FLAG (C609S) | R1c609sN_fwd | tttctgacttatagtcgctttgttaaatcatATGGATTCATTGTCTGCATATC |
| | R1c609sN_rev | caaacaaggatctagACTAACACTACAGTTGAAATCC |
| | R1c609sC_fwd | aactgtagtgttagtCTAGATCCTTGTTTGGCTC |
| | R1c609sC_rev | tagtctttatcatcgtcgtccttgtagtcggatccTGAATTAAGCCCAAATAGATATAG |
| Rrp2-FLAG | Rrp2_fwd | caactaattattcgaaacggaattcgaaacgATGAGAAATAATACAGCTTTTGAAC |
| | Rrp2_rev | tttaaatggccggccggtaccTTATCGTGATGACATTCCAAATAAAAATG |
| Rrp2-DAEA-FLAG (D539A; E540A) | Rrp2D539A_fwd | caactaattattcgaaacggaattcgaaacgATGAGAAATAATACAGCTTTTGAAC |
| | Rrp2D539A_rev | tttgagcggcCGCCAATATAACTCGATACC |
| | Rrp2E540A_fwd | tatattggcgGCCGCTCAAACTATCAAAAAC |
| | Rrp2E540A_rev | tttaaatggccggccggtaccTTATCGTGATGACATTCCAAATAAAAATG |
| Rrp2-CS-FLAG (C749S) | R2Nc749s_fwd | tttctgacttatagtcgctttgttaaatcatATGAGAAATAATACAGCTTTTGAAC |
| | R2Nc749s_rev | tgcaacaacatccatGGATAAGGAGCATTGCAAC |
| | R2Cc749s_fwd | caatgctccttatccATGGATGTTGTTGCAGAAC |
| | R2Cc749s_rev | tagtctttatcatcgtcgtccttgtagtcggatccTCGTGATGACATTCCAAATAAAAATG |
| Primers used for qPCR in ChIP experiments | | |
| *act1* | act_F | cgccgaacgtgaaattgttcgtga |
| | act_R | aagggaggaagattgagcagcagt |
| centromere *cnt* | cnt_F | caaccgttgcaacttacatcagca |
| | cnt_R | ccggtcgccaaatagcaatgagat |
| centromere *dg* | dg_F | taccgtgattagccttactccgca |
| | dg_R | accgcaagatagagtaggatgggt |
| telomere | tel_F | tcaaagttggcgacgttgctgatg |
| | tel_R | aagcaatgtgtggagcaacagtgg |

**Suplementary Figures**

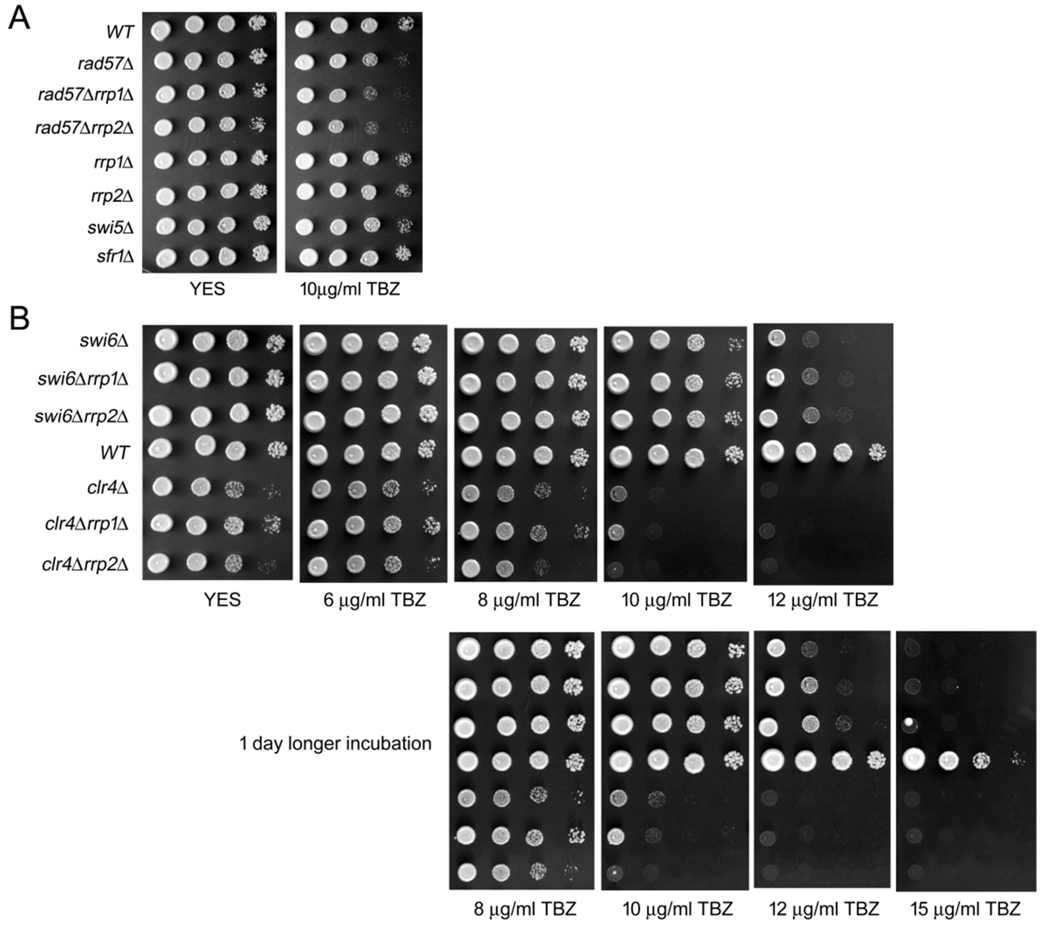


**Fig. S1 Rrp1 and Rrp2 may be implicated in centromere function**
(A) Strains devoid of genes for Swi5-Sfr1 mediator complex and Rrp1 or Rrp2 are not sensitive to TBZ. *rad57*Δ is sensitive and deletion of *rrp1*+ or *rrp2*+ in this mutant increases this sensitivity. (B) Deletion of *rrp1*+ or *rrp2*+ decreases TBZ sensitivity in *swi6*Δ mutant while in *clr4*Δ mutant, deletion of *rrp1*+ but not *rrp2*+ rescues its growth defect.

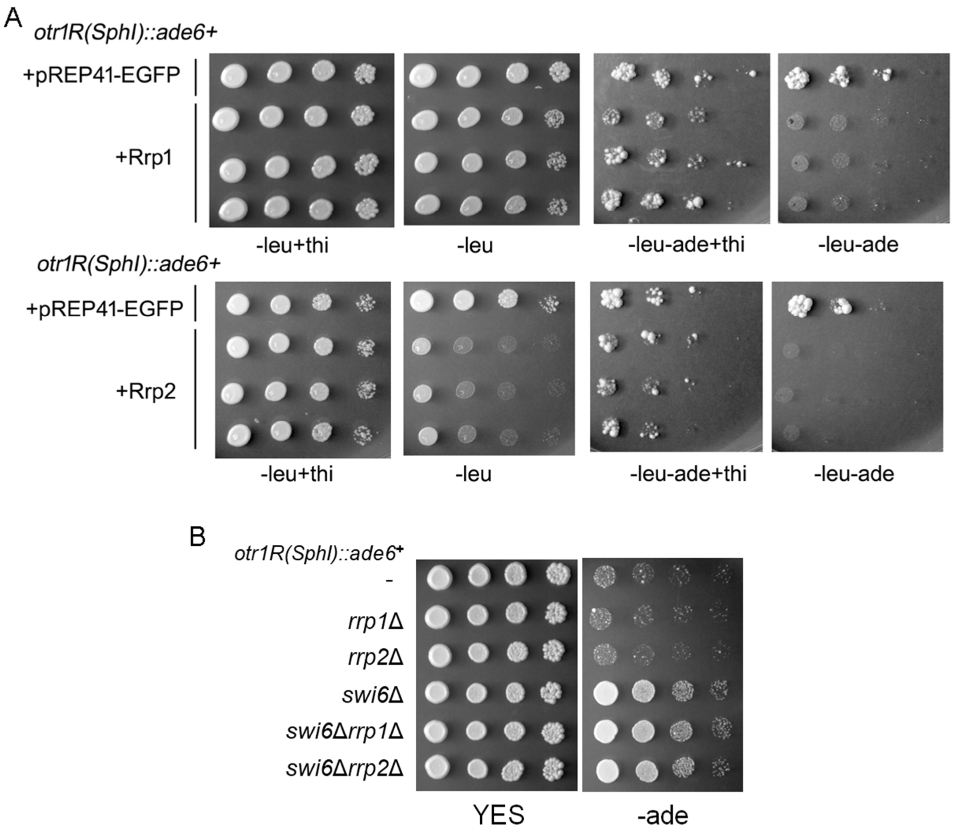


**Fig. S2 The effect of Rrp1 and Rrp2 on transcriptional silencing at the centromere**
(A) *rrp1+* or (B) *rrp2+* over-expression results in an increase in repression of the *dg-ade6+* gene as determined by the lack of growth on plates devoid of adenine and thiamine (expression inducing conditions) (B) Deletion of *rrp1+* or *rrp2+* has no effect on the silencing of *dg-ade6+* gene in *swi6+* or *swi6*Δ backgrounds, monitored by the ability of these mutants to grow on plates without adenine.

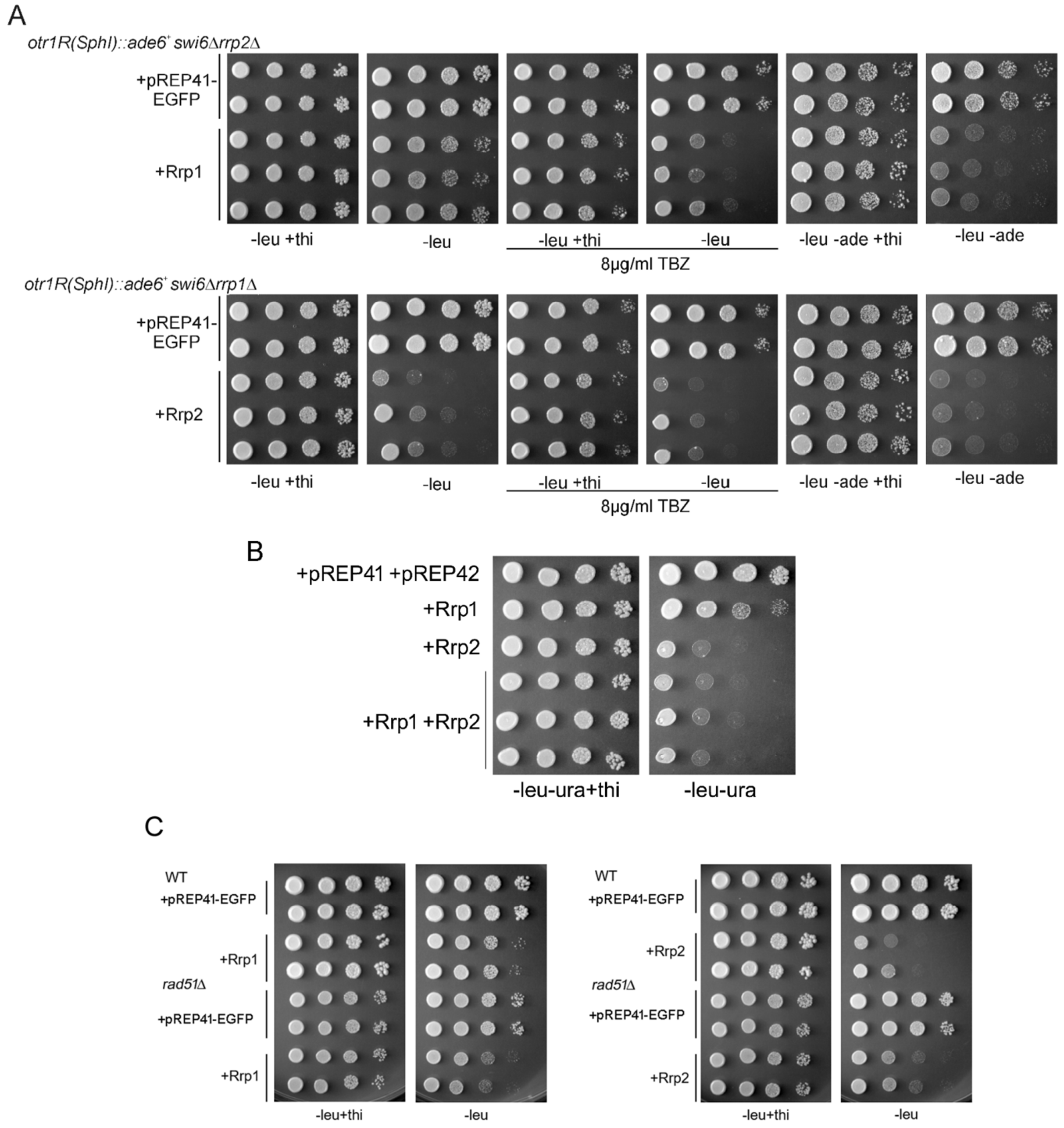


**Fig. S3 Interdependence of *rrp1*+ and *rrp2*+ over-expression phenotypes**
(A) Centromere functions of Rrp1 and Rrp2 are not dependent on the presence of the other respective paralogue. Reporter *swi6*Δ*dg-ade6*+ strain with *rrp1*+ or *rrp2*+ deletion was transformed with pREP41-EGFP-Rrp2 or pREP41-EGFP-Rrp1 plasmid, respectively. Loss of viability and increased TBZ sensitivity are observed as growth inhibition while transcriptional repression by the ability to grow on plates lacking adenine under expression induction conditions (plates lacking thiamine). (B) Growth defect resulting from simultaneous over-expression of both genes is similar to the effect caused by *rrp2*+ over-expression. Strains were transformed with empty pREP41-EGFP or pREP42-EGFP vector (carrying respectively leucine or uracil marker gene) as control and corresponding plasmids with *rrp1*+ and *rrp2*+ genes. Loss of viability is observed as growth inhibition on plates lacking thiamine (expression induction conditions). (C) Only growth defect resulting from *rrp2*+ over-expression is dependent on the presence of Rad51 recombinase.

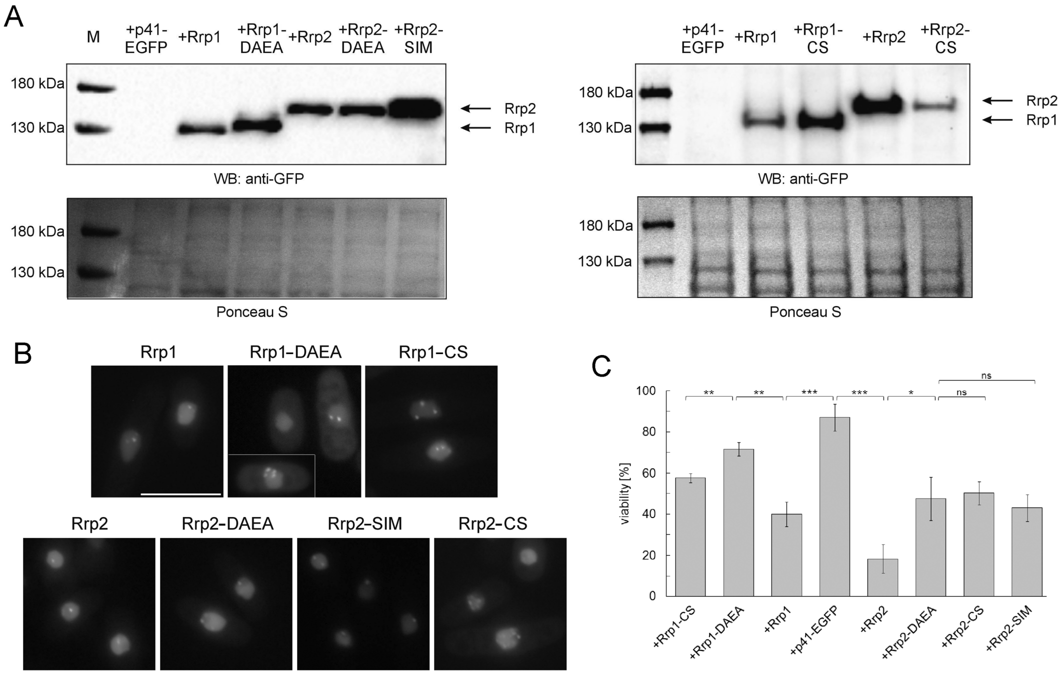


**Fig. S4 Analysis of the functionality of Rrp1 and Rrp2 mutations in ATPase Walker-B domain (Rrp1-DAEA, Rrp2-DAEA), ubiquitin ligase RING domain (Rrp1-CS, Rrp2-CS) and SUMO interacting motif (Rrp2-SIM)**

WT strain over-expressing wild type or mutated versions of *rrp1*+ or *rrp2*+ from pREP41-EGFP plasmid was used. (A) All proteins are expressed as confirmed by Western blot with anti-GFP antibodies of proteins isolated from cultures incubated for 24 h without thiamine (expression inducing conditions). (B) All proteins form foci in the nucleus. (C) All domains contribute to the loss of viability conferred by *rrp1*+ or *rrp2*+ over-expression. Loss of viability was determined for strains growing in the presence and absence of thiamine, as compared to empty vector pREP41-EGFP control. The experiment was repeated at least twice for two independent transformants. The error bars represent the standard deviation about the mean values. Student's t-test was used to calculate the P-value (* $0.01 < \text{P-value} \leq 0.05$, ** $0.001 < \text{P-value} \leq 0.01$, *** $\text{P-value} \leq 0.001$). Cells from cultures used in (A) were analysed by fluorescence microscopy (B). Scale bar indicates 10 µm.

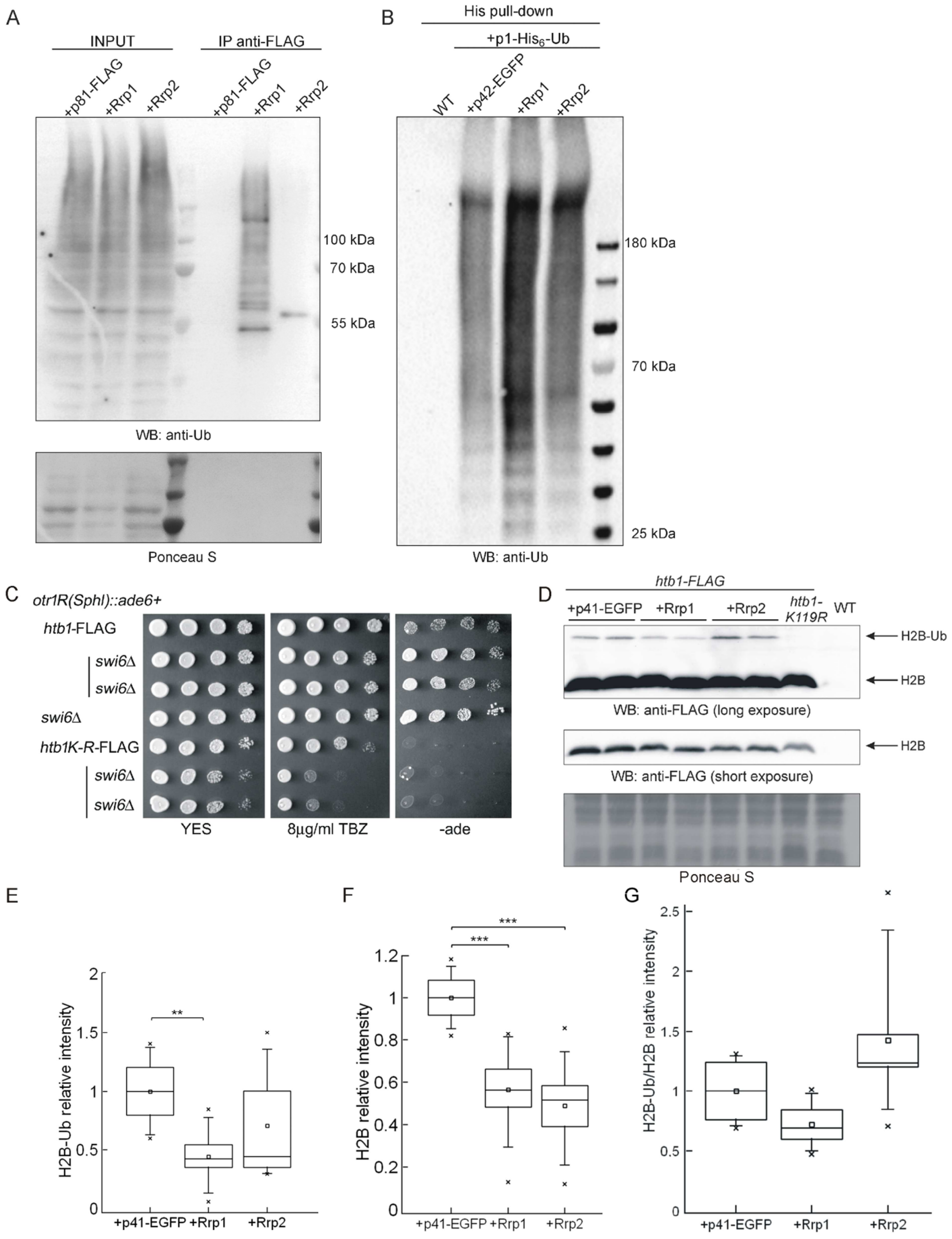


**Fig. S5 Rrp1 and Rrp2 are potential ubiquitin ligases that can affect histone density but their role in centromere function does not involve histone H2B-K119 ubiquitylation**
(A) Rrp1 and Rrp2 may form complex with ubiquitylated proteins. After 24 h growth in minimal medium in the absence of thiamine (expression inducing conditions) proteins were immunoprecipitated with anti-FLAG agarose (ANTI-FLAG® M2 Affinity Gel) from strains transformed with pREP81-Rrp1-FLAG, pREP81-Rrp2-FLAG plasmids and with empty vector as control and subjected to Western blot analysis. Ubiquitylated with proteins were detected with anti-ubiquitin antibody. (B) Rrp1 and Rrp2 overproduction leads to accumulation of ubiquitin modified proteins. WT strain was co-transformed with pREP1-His$_6$-Ub plasmid

together with pREP42-EGFP-Rrp1, pREP42-EGFP-Rrp2 or empty vector as control. After 24 h of growth in minimal medium in the absence of thiamine (expression inducing conditions) ubiquitylated proteins were isolated by His pull-down and detected at the Western blot with anti-ubiquitin antibody. (C) *H2B-K119R* mutation, as *rrp1*+ or *rrp2*+ over-expression, strongly sensitizes *swi6*Δ cells to TBZ and reverses the transcriptional de-repression of the *dg-ade6*+ gene conferred by *swi6*Δ. (D) Rrp1 or Rrp2 have no effect on H2B-K119 ubiquitylation, but affect unmodified histone H2B levels. Total protein extracts were prepared from *htb1*-FLAG strain over-expressing *rrp1*+ or *rrp2*+ genes, and probed with anti-FLAG antibodies. A mutant *htb1-K119R*-FLAG strain was used as control to identify a band corresponding to ubiquitylated H2B (H2B-Ub). Data were quantified and shown as box and whiskers plots representing (E) relative intensity of H2B-Ub versus Ponceau S loading control, (F) relative intensity of H2B signal versus Ponceau S loading control and (G) relative intensity of H2B-Ub versus H2B signal, all reads were normalised by mean value obtained for vector control samples. Western blots were analysed by ImageLab Software and Ponceau S stained membranes by ImageJ Software. Three independent Western blots from three separate protein isolations from two different transformants were examined. Box represents the range from 25 to 75%, whiskers—the range from 5 to 95%, line dividing the box—the median and a full square—the mean value. Student's t-test was used to calculate the P-value (* 0.01 < P-value ≤ 0.05, ** 0.001 < P-value ≤ 0.01, *** P-value ≤ 0.001).

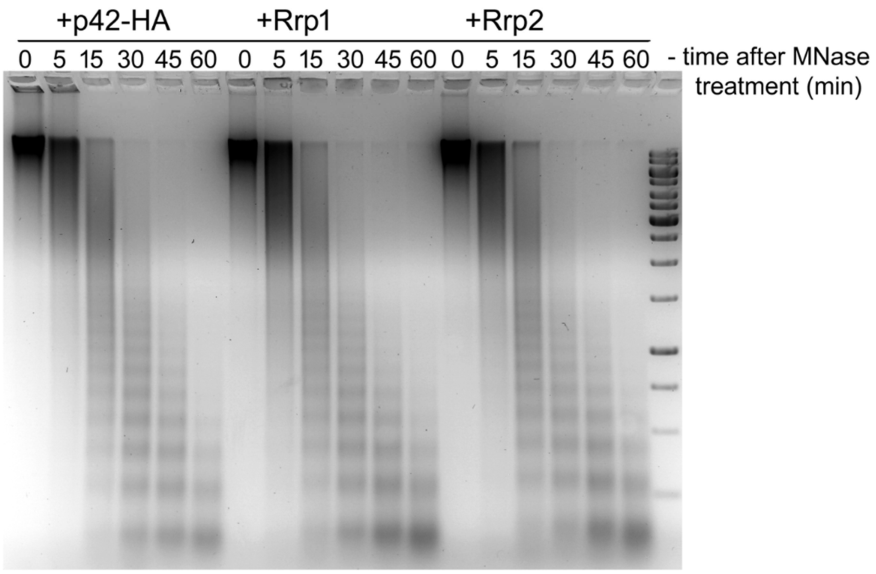


**Fig. S6 Global nucleosome spacing is not altered by Rrp1 or Rrp2 overproduction**. MNase assay of chromatin extracts from cells transformed with plasmids carrying genes for Rrp1-HA or Rrp2-HA and. Simply safe stained gel with partially digested chromatin samples treated for different times with 5 U/ml MNase. The assay was repeated twice.
